# Structural basis for alternative 3′ splice site selection in the human spliceosome active center

**DOI:** 10.64898/2026.08.05.741260

**Authors:** Gabriele Marciano, Simon Eckert, Luíza Zuvanov, Takero Miyagawa, Leo Yang, George-Valentin Datcu, Yuewen Sheng, Hye Young Kwon, Louis Cameron, Marco Preußner, Florian Heyd, Sebastian M. Fica

## Abstract

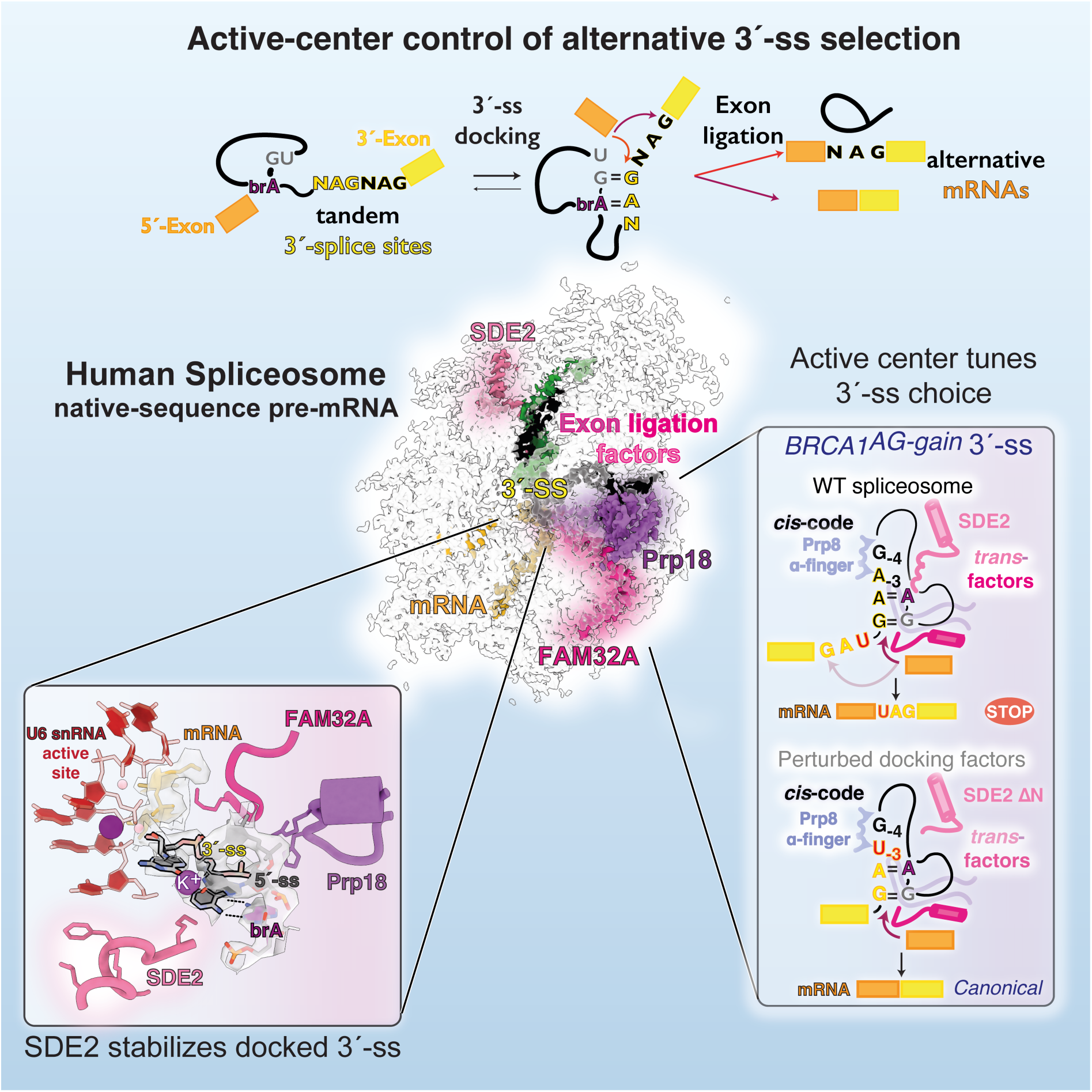

**Key findings:**

- Cryo-EM reveals SDE2 as a novel factor stabilizing the spliceosome active center
- SDE2 ΔN structures visualize a stalled C* state with impaired docking-factor engagement.
- SDE2, Prp18, and FAM32A read a cis-code to stabilize weaker proximal 3′-ss.
- SDE2 ΔN destabilizes 3′-ss docking and rescues *BRCA1* and *CFTR* mis-splicing *in vivo*.

Accurate alternative splicing requires discrimination between adjacent 3′ splice sites (3′-ss) during catalysis and is disrupted by pathogenic AG-gain mutations that create competing 3′-ss. Here, we present cryo-EM structures of human spliceosomes assembled on native-sequence pre-mRNAs, revealing how the catalytic core controls alternative 3′-ss selection. SDE2 is a previously unrecognized active-center component that promotes a docking-competent spliceosome conformation. Machine learning, *in vivo* transcriptomics, and *in vitro* biochemistry show how SDE2 cooperates with FAM32A and Prp18 to act as readers of a cis-regulatory code that governs 3′-ss selection during catalysis. These factors promote weaker, proximal site use by counteracting an intrinsic distal bias generated by active-site interactions with the distal-site −4 nucleotide. Structural or genetic perturbation of these exon-ligation factors destabilizes proximal 3′-ss docking and restores canonical splicing in disease-relevant *CFTR* and *BRCA1* AG-gain alleles. Our work establishes the spliceosome active center as a tunable regulatory hub for alternative splicing.

## Introduction

Alternative splice-site selection expands proteome diversity^1^ and regulates gene expression^2^. Because accurate and efficient intron removal determines transcript fate and produces regulatory RNAs^3–6^, the spliceosome must precisely balance fidelity with plasticity. To excise introns from pre-mRNAs and ligate exons to form mature transcripts, the spliceosome uses conserved elements, including the 5′ splice site (5′-ss), the branch point (BP), and the 3′ splice site (3′-ss), to define intron boundaries (Figure 1A).

**Figure 1.**
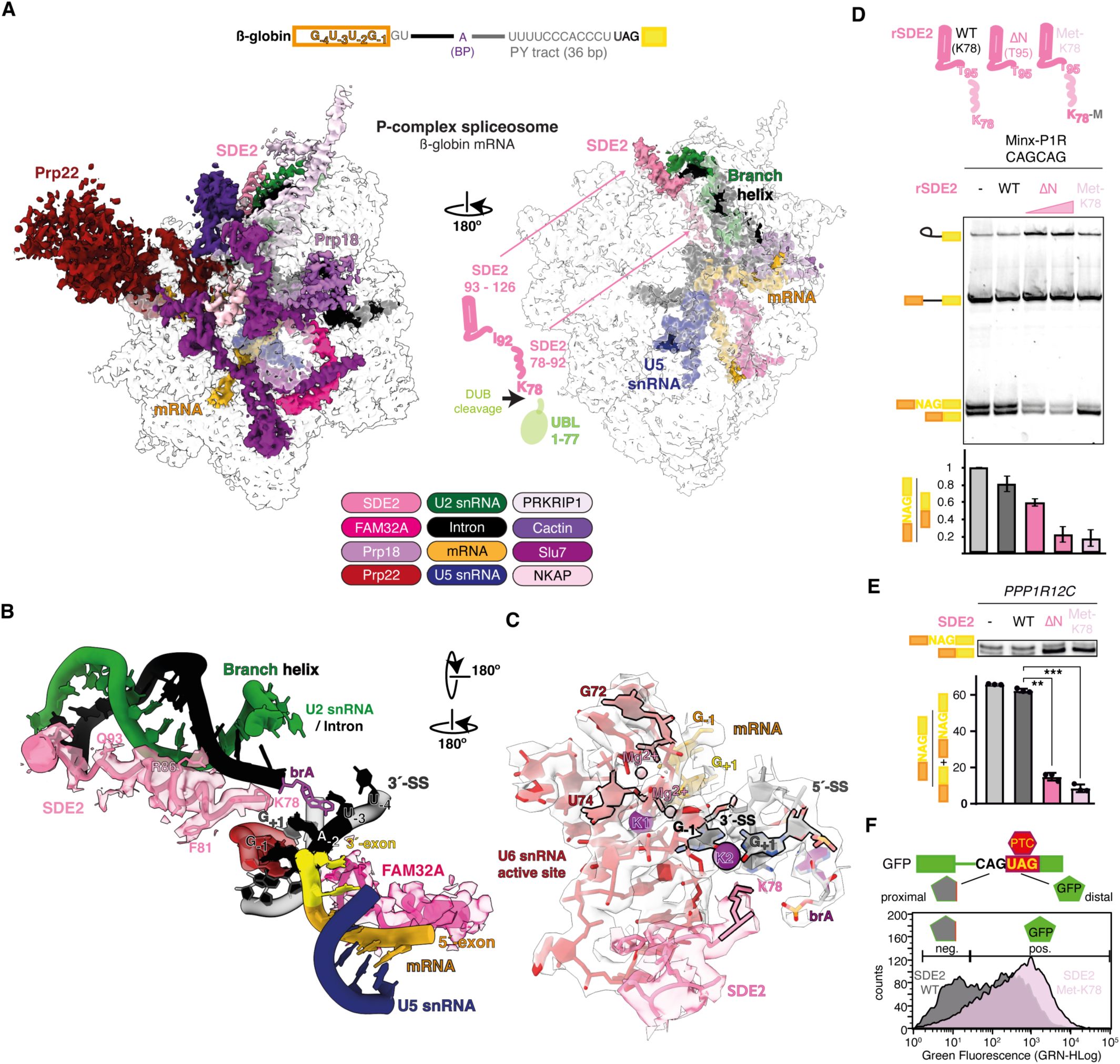
SDE2 is a novel active-center component that modulates alternative 3′-ss selection. **(A)** Architecture of the β-globin P complex, highlighting SDE2 and the 3′-ss docking network. **(B)** Exon-ligation factors SDE2 and FAM32A clamp the docked 3′-ss. **(C)** U6 snRNA active site, showing SDE2 K78 near the K⁺-mediated pairing of the docked 3′-ss and 5′-ss. **(D)** *In vitro* splicing. Recombinant SDE2 ΔN impairs overall exon ligation and shifts usage to the distal 3′-ss. The Met-K78 mutant maintains exon-ligation efficiency but acts purely as a regulatory switch, shifting 3′-ss use. Data are represented as mean ± SD (n = 4). **(E)** *In vivo* RT–PCR of a *PPP1R12C* minigene confirms that SDE2 mutations promote distal 3′-ss usage. Data are represented as mean ± SD (n = 3); **, p < 0.01; ***, p < 0.001. **(F)** SDE2 disruption shifts NAGNAG selection in a GFP reporter *in vivo*, activating distal GFP expression. Representative FACS is shown (quantification in Figure S2).

While early splice-site recognition is increasingly well understood^7–9^, the regulation of tandem alternative 3′-ss (TASSs) remains poorly defined. For TASSs, in humans typically NAGNAG motifs, the spliceosome must discriminate between two closely spaced 3′-ss AGs separated by only three nucleotides^10^. This local 3′-ss plasticity is conserved across vertebrates and promotes the evolution of tissue-specific proteome diversity^10–13^, but creates functional vulnerability. *De novo* AG-gain mutations between the BP and canonical 3′-ss, as well as point mutations in existing TASSs, drive pathogenic mis-splicing in diverse human diseases^14,15^.

Current models of alternative splicing primarily attribute splice-site choice to early assembly events mediated by factors such as U2AF, which defines the 3′-ss and controls usage of cassette exons^8,16^. However, discrimination between TASSs requires the spliceosome to choose between adjacent AGs during the catalytic stage. Cryo-EM structures of yeast and human spliceosomes have delineated the extensive conformational remodeling during transition from the C to the C* complex, when recruitment of specific factors like FAM32A promotes pairing of the 3′-ss and 5′-ss for active-site docking and renders the C* spliceosome competent for exon ligation^17–24^. Indeed, recent functional screens indicate that specific exon-ligation factors, including FAM32A and SDE2, modulate NAGNAG usage^25^. However, all previously reported human C* and P spliceosome structures were determined using viral-derived pre-mRNAs with optimal splice sites and did not visualize several key exon-ligation factors, including Prp18, engaged with substrate RNA^18,24–27^. Consequently, the molecular and structural mechanisms through which the active site accommodates weaker, native 3′-ss and discriminates between competing TASSs during catalysis remain obscure.

Here we determine cryo-EM structures of the human spliceosome assembled on native intron sequences to define the mechanism of tandem 3′-ss selection during exon ligation. The structures reveal SDE2 as a previously unrecognized active-center component that stabilizes a docking-competent C* conformation and promotes catalytic progression. *In vitro* and *in vivo* assays, combined with machine learning, demonstrate how SDE2, together with Prp18 and FAM32A, facilitates docking of weaker proximal sites against an intrinsic, sequence-encoded bias towards stronger distal sites. By applying this mechanistic understanding to functional *in vivo* assays, we demonstrate the potential to correct mis-splicing of disease-associated NAGNAG variants. Together, our results define a comprehensive molecular framework for alternative 3′-ss selection during catalysis and identify active-site docking as a potential future target for therapeutic intervention.

## Results

### SDE2 is a novel active-center component that governs alternative 3′-ss choice

To visualize how the human spliceosome accommodates native 3′-ss sequences, we determined a 3.2–3.5 Å cryo-EM structure of a β-globin P complex stalled immediately after exon ligation (Figure 1A, Figure S1, A to C, Tables 1 and S1). The reconstruction reveals the complete architecture of the human exon-ligation active site, including clear density for the docked 3′ splice site (3′-ss) and associated mRNA (Figure S1, D to J). Notably, the structure resolves SDE2 acting cooperatively with Prp18 and FAM32A in the catalytic center (Figure 1A and B), where they form a docking network that stabilizes 3′-ss engagement.

**Table 1.**
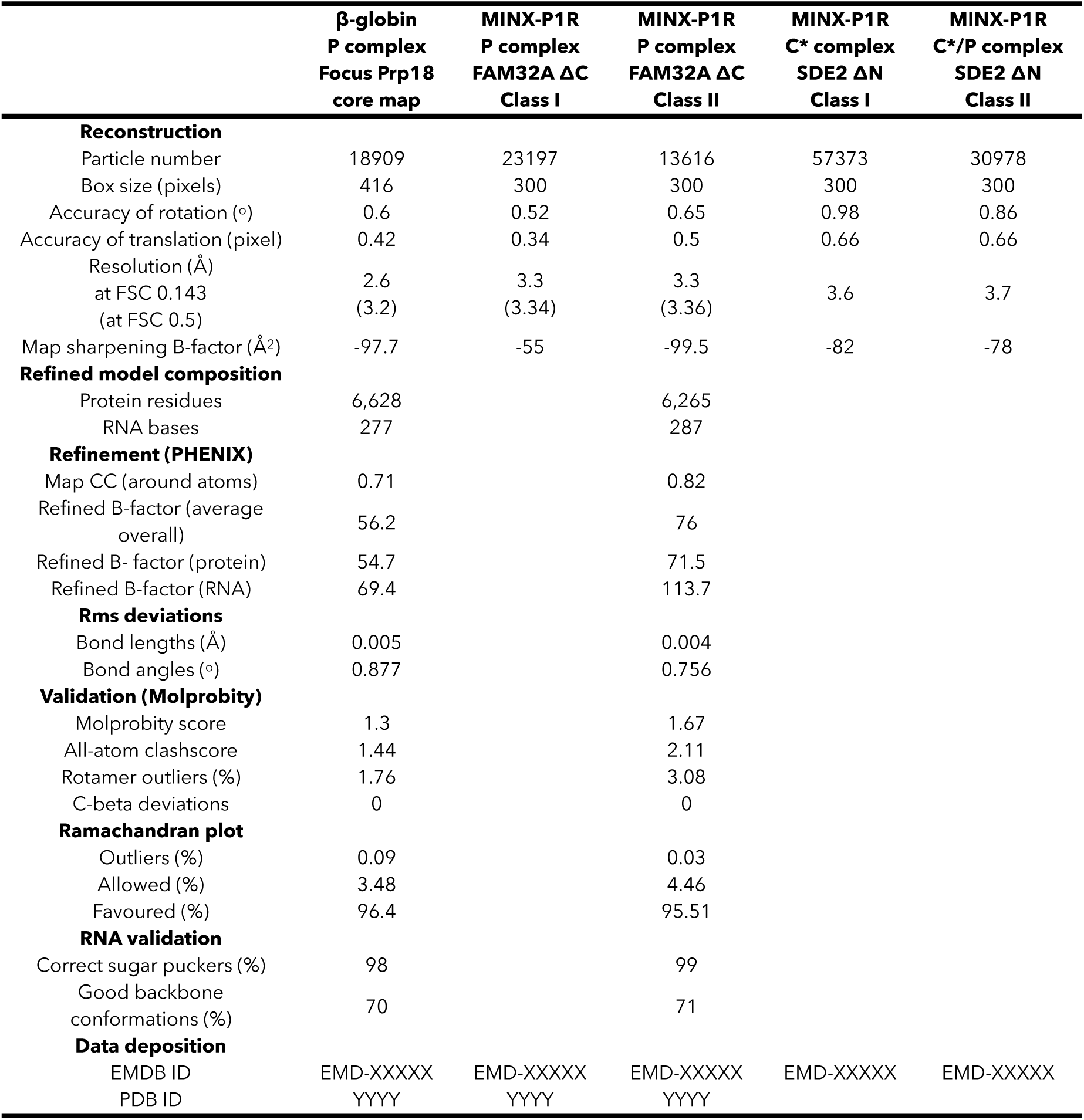
Cryo-EM refinement statistics. Data collection and refinement parameters are shown for the individual final maps used for model building and structural analysis.

SDE2 binds the human spliceosome during exon ligation^24^, following cleavage of its N-terminal UBL domain^28,29^. While residues immediately downstream of this cleavage site were invisible in cryo-EM maps assembled on viral pre-mRNAs, our β-globin map reveals ordered density for the SDE2 N-terminus (residues 78–104), which runs along the branch helix and extends deep into the active center (Figure 1, B and C). Here, SDE2 contacts the U2 snRNA, U6 snRNA, the branch adenosine, and the intron to precisely position the 3′-ss for exon ligation (Figure 1, B and C). Our β-globin structure thus reveals SDE2 as a previously unrecognized core component of the spliceosome active center. SDE2 stabilizes the active-site architecture required to position the docked 3′-ss leaving group for nucleophilic attack, and cooperates with FAM32A, which clamps the upstream 5′-exon onto U5 snRNA (Figure 1B), to align both reactants for exon ligation.

Within the active-center, the exposed N-terminal SDE2 residue K78 plays a key role in 3′-ss docking. The cryo-EM density places K78 in multiple conformations adjacent to the docked 3′-ss, consistent with a role in supporting K⁺-mediated pairing between the 5′-ss and 3′-ss (Figure 1C, c.f. ^22^). Thus K78 may stabilize the local RNA backbone for 3′-ss docking by helping to organize a water-mediated coordination environment at the K2 site, in a manner reminiscent of lysine-stabilized RNA pairing in the *lysC* riboswitch^30^ and of basic residues that organize water-mediated K⁺ pockets within the ribosome^31^. To test the function of the N-terminal K78 during 3′-ss selection, we inserted a methionine upstream of K78 (Met-K78, Figure S2 A and B). Notably, this precise methionine insertion did not affect exon-ligation efficiency *in vitro* and did not accumulate pre-mRNA *in vivo* (Figure 1D; Figure S2, C to D). Instead, the SDE2 Met-K78 mutant acted as a regulatory switch, shifting splice-site usage toward distal 3′-ss acceptors *in vitro* and for endogenous introns *in vivo* (Figure 1, D and E; Figure S2, D and E). Perturbation of this single SDE2 residue also increased distal site usage and GFP expression in a CAGUAG reporter, by skipping inclusion of the distal site stop codon, thus demonstrating the functional relevance of TASS regulation for protein production *in vivo* (Figure 1F; Figure S2, F and G).

While K78 acts as a regulatory switch for TASS selection, the strict evolutionary conservation of the broader SDE2 N-terminal helix and its extensive contacts with the branch helix (Figure 1B; Figure S2, H and K) suggested an additional function. Indeed, deleting the entire helical SDE2 N-terminus (ΔN) severely impaired overall exon-ligation efficiency and reduced RNA binding *in vitro* (Figure 1D, Figure S2,I and J). Supporting a structural role in stabilizing the branch helix during C-to-C* remodeling, the SDE2 N-terminus is only ordered in our β-globin structure, which features a weaker branch helix compared to spliceosomes assembled on the viral Minx pre-mRNA (Figure S2K). Thus without the SDE2 N-terminus spliceosomes cannot efficiently progress through the splicing pathway. Indeed, deleting the SDE2 N-terminus caused accumulation of lariat intermediates in pre-catalytic C*-like complexes in glycerol gradient sedimentation assays (Figure S3A). *In vivo*, this defect manifested as the selective accumulation of pre-mRNA (intron retention) for specific minigenes (Figure S2L), as observed previously following impairment of other exon-ligation factors^32^. Because deletion of the N-terminus inherently removes the K78 regulatory residue, SDE2 ΔN also shifted splicing towards distal NAGNAG acceptors *in vitro* and *in vivo* (Figure 1,D and E, Figure S2,D,E, and L). Together, these data indicate that the SDE2 N-terminus has dual functions: its terminal K78 tunes alternative 3′-ss choice, while the extended N-terminal helix stabilizes the exon-ligation conformation, thus ensuring efficient catalytic progression.

### SDE2 stabilizes the C* conformation to facilitate 3′-ss selection with FAM32A and Prp18

In our β-globin structure, engagement of SDE2 correlates with strong density for additional exon-ligation factors, notably Prp18 and FAM32A (Figure 2A and B Figures S1F and S2K). To define the molecular pathway for recruiting this network of exon-ligation factors, we determined cryo-EM structures of spliceosomes assembled in the presence of the SDE2 ΔN mutant (Figure 2C; Figure S3, B to E). Remarkably, 3D classification revealed that without the SDE2 N-terminus, the majority of spliceosomes (66%, Class I) accumulate in a C*-like conformation where key exon-ligation factors, including FAM32A, Prp22, and Slu7, fail to stably engage the active center. Only a minority of complexes (33%, Class II) showed stable engagement of exon-ligation factors in a complete P-like state (Figure 2C, Figure S3F), consistent with the accumulation of lariat intermediates in pre-catalytic C*-like complexes in glycerol gradient sedimentation assays (Figure S3A). In the absence of the SDE2 N-terminus, the spliceosome population effectively shifts away from a factor-engaged state, leading to a poorly docked 3′-ss (Figure S3F). Supporting a cooperative role in stabilizing this C* docked state, deletion of the FAM32A C-terminus (FAM32A ΔC) phenocopied the distal splice-site switch caused by SDE2 loss without impairing exon ligation on its own, but exacerbated the exon-ligation defects caused by SDE2 depletion or mutation (Figure 2D; Figure S4, A to C). Indeed, deletion of the SDE2 N-terminus strongly reduced the kinetics of exon ligation at a tandem NAGNAG 3′-ss (Figure S4, D to F). Our data suggest a model in which SDE2 promotes a docking-competent C* conformation and facilitates efficient recruitment of additional exon-ligation factors, thereby stabilizing the spliceosome catalytic center for exon ligation (see also Supplementary Text).

**Figure 2.**
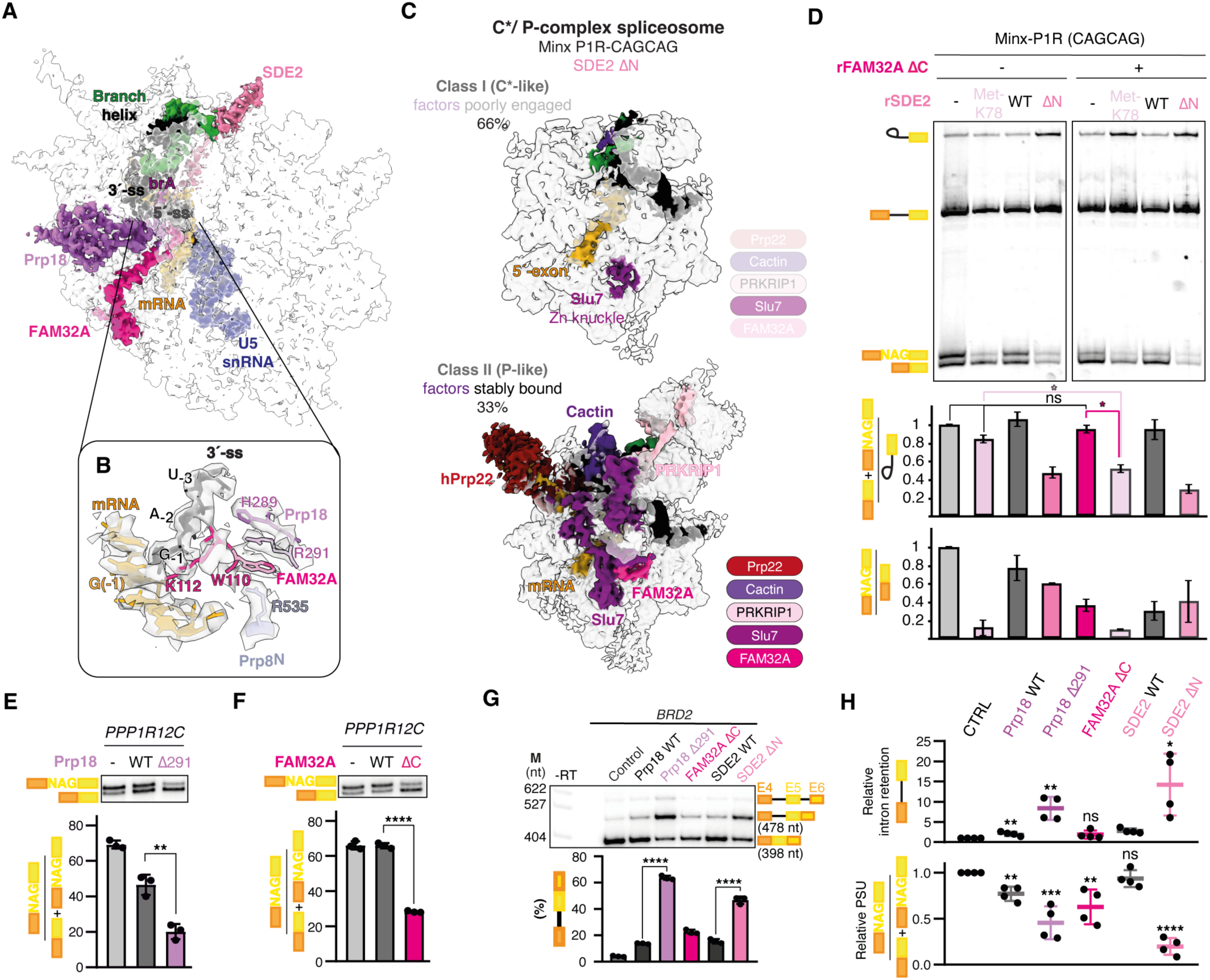
SDE2 promotes engagement of exon-ligation factors in a docking-competent C* conformation to facilitate catalytic progression and alternative 3′-ss use. **(A and B)** Cryo-EM structure of the β-globin P complex showing Prp18, SDE2, and FAM32A engaging the catalytic core **(A)** and stabilizing the docked 3′-ss **(B)**. **(C)** The SDE2 N-terminus stabilizes a catalytically competent C* conformation. Major cryo-EM classes for spliceosomes assembled with SDE2 ΔN show either poorly engaged (Class I) or stably bound (Class II) exon-ligation factors. (D) *In vitro* splicing of a Minx-P1R CAGCAG substrate demonstrates that FAM32A ΔC and SDE2 mutants shift splicing to the distal 3′-ss. **(E and F)** Prp18 Δ291 **(E)** and FAM32A ΔC **(F)** decrease proximal NAGNAG 3′-ss usage *in vivo* on a *PPP1R12C* minigene. Data are represented as mean ± SD (n = 3). **(G)** Prp18 and SDE2 mutants cause strong, intron-specific pre-mRNA retention at the endogenous *BRD2* locus. Data are represented as mean ± SD (n = 3). **(H)** Across multiple human heterologous minigenes, perturbation of Prp18 and SDE2 causes increased intron retention (top). Perturbations of FAM32A, Prp18, and SDE2 have comparable effects on NAGNAG choice. Data are represented as mean ± SD for 4 independent minigenes, with each individual point representing the average of 3 replicates for a specific minigene. In all panels, statistical significance by unpaired t-test: *, p < 0.05; **, p < 0.01; ***, p < 0.001; ****, p < 0.0001; ns, not significant.

In addition to revealing the fundamental role of the SDE2 N-terminus, our β-globin structure provides crucial insights into how the FAM32A-Prp18-SDE2 network selects 3′-ss during exon ligation. Remarkably, this is the first human spliceosome structure to visualize Prp18 (PRPF18) (Figure 2A), which had previously been observed only in yeast spliceosome structures, where it promotes 3′-ss fidelity^33,34^. Human Prp18 directly promotes β-globin splicing^35^ (Figure S5,A to F) by binding the Prp8 RNase H domain, while its conserved loop abuts the 3′-ss to create a dedicated channel for 3′-ss docking (Figure S5C). Our maps also reveal that Prp18 R291 binds and stabilizes the C-terminus of FAM32A, while H289 bridges this interaction with the phosphate backbone of the docked 3′-ss (Figure 2B; Figure S5, G and H). Consistent with a crucial role of Prp18 R291 in active site geometry, its mutation forced distal 3′-ss usage *in vitro* and its deletion shifted usage to distal sites across diverse native substrates *in vivo* (Figure 2E; Figure S5I and S6A), precisely mirroring the effects of deleting the FAM32A C-terminus (Figure 2F; Figure S5I and Figure S6B) or the SDE2 N-terminus (Figure S2L). *In vitro* depletion and add-back assays further confirmed that FAM32A regulates proximal 3′-ss selection during the C to C* transition (Figure S6, C to G). Importantly, the alternative splicing events regulated by FAM32A and Prp18 were insensitive to knock-down of U2AF (Figure S6, H to K), demonstrating that TASS selection by exon-ligation factors is mechanistically distinct from the early spliceosome assembly steps, when U2AF regulates alternative cassette exons.

Transcriptome profiling extended these mechanistic conclusions genome-wide, defining the dual functions of this C* docking network. First, specific exon-ligation factors are required for catalytic progression through the splicing pathway. Over-expressing exon-ligation factor mutants (Prp18 Δ291 and SDE2 ΔN) caused selective intron retention in model minigenes and at sensitive endogenous loci (Figure 2G and H; Figure S7, A to D). Second, for transcripts that successfully reach the C* stage, the 3′-ss docking network acts as a regulatory hub. All perturbations of the C* network factors altered TASS usage, unidirectionally shifting NAGNAG events towards the distal site (Figure 2H; Figure S7A, E and F). Thus FAM32A, Prp18, and SDE2 form an integrated active-site docking network that ensures efficient spliceosome progression while governing the precision of tandem 3′-ss choice during catalysis.

### Active-center geometry establishes a cis-regulatory code for 3′-ss docking

SDE2, Prp18, and FAM32A provide a *trans*-acting network that stabilizes proximal 3′-ss. However, local pre-mRNA *cis*-features also impact TASS choice^11,36^ (Figure S7, G to K). Our β-globin P complex structure reveals the underlying molecular mechanism for this sequence-specific bias. In the docked conformation, the 3′-ss stacks coaxially onto the U6/5′-ss helix, forming an extended RNA stack capped by the Prp8 α-finger (Figure 3, A and B). Here, we observe for the first time how Prp8 F1509 stacks against the −4 nucleotide of the 3′-ss and introduces a backbone bend that guides the AG dinucleotide into the active site (Figure 3B). Because aromatic stacking is favored by purines, and is strongest for guanosine^37^, this architecture predicts a sequence bias that favors sites with a −4 purine, whereas a suboptimal −4 nucleotide would increase dependence on exon-ligation factors (Figure 3B). The distal site of a tandem NAGNAG invariably contains an optimal −4G, and thus possesses high intrinsic docking stability, creating an inherent distal bias. Consistent with this mechanism, mutating a weaker-stacking −4 cytosine at a proximal 3′-ss to a guanosine (−4G; GCAGCAG) was sufficient to drive near-complete proximal usage *in vitro* and *in vivo* (Figure 3, C to F), and rendered the 3′-ss poorly responsive to mutation of exon-ligation factors (Figure 3, C to F). Together with known Prp8 α-finger interactions with the −3 position^21,24,38^, our newly discovered −4 interaction provides a structural basis for the *cis*-code of NAGNAG selection and completes the mechanism for guiding the 3′-ss into the active site.

**Figure 3.**
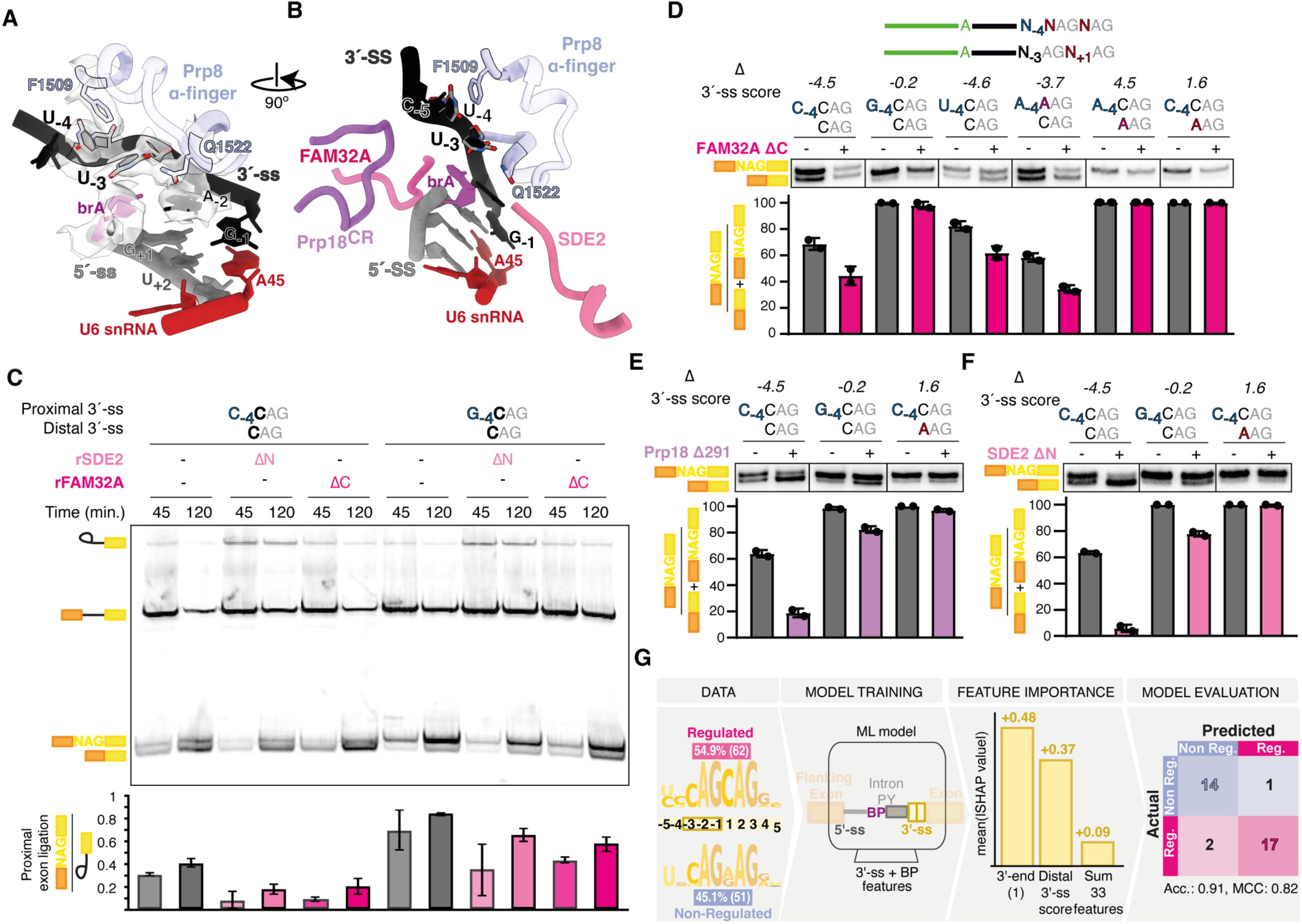
Active-center geometry and a cis-regulatory code explain how exon-ligation factors govern alternative 3′-ss use. **>(A and B)** Structure of the docked 3′-ss in the β-globin P-complex active site, showing stacking onto the U6/5′-ss helix and capping by the Prp8 α-finger. Prp8 F1509 (not Prp18) stacks on the −4 nucleotide of the 3′-ss **(A)**, while exon-ligation factors bend and contact the 3′-ss backbone **(B)**. **(C)** The −4 nucleotide at the 3′-ss controls docking and 3′-ss selection, independently of exon-ligation factors. *In vitro* splicing assays using Minx-P1R substrates with indicated proximal 3′-ss −4 position variants (C-4 vs. G-4). Data are represented as mean ± SD (n = 3). **(D to F)** Validation of model-predicted *cis*-determinants by minigene assays *in vivo* upon expression of FAM32A ΔC **(D)**, Prp18 Δ291 **(E)**, and SDE2 ΔN **(F)**. Sequence cartoons indicate the local NAGNAG context and corresponding Δ 3′-ss scores. Representative gels (middle) and quantification of proximal vs. distal isoform usage (bottom) are shown. Data are represented as mean ± SD (n = 2). **(G)** Machine learning (ML) framework defining the *cis*-regulatory code for alternative 3′-ss selection. The model integrates sequence motifs of regulated versus non-regulated targets (left) to identify predictive *cis*-features, revealing via SHAP value analysis that the 3′-end nucleotide and distal 3′-ss score are the primary determinants of regulatory dependency (center), yielding a highly accurate predictive model (confusion matrix, right).

To test whether this structural code governs NAGNAG selection across the transcriptome, we trained a machine-learning (ML) model on FAM32A-regulated versus non-regulated introns^25^ (Figure 3G). The model identified distal 3′-ss strength as the dominant predictive determinant of regulation by C* docking-network components. FAM32A target introns are strongly enriched for a strong distal CAG motif and have a weak proximal −4 position (C/U) (Figure 3G, Figure S8A-F). Weakening distal docking elements (suboptimal CAGAAG) reduced factor dependence, whereas weakening the proximal site amplified the proximal-distal *cis*-score difference, rendering the proximal site highly dependent on the docking network *in vivo* (Figure 3D-F; Figure S8G). To visualize the physical basis for this functional dependency, we determined cryo-EM structures of P-spliceosomes assembled with the FAM32A ΔC mutant, which would disrupt the docking network (Figure S9, A to G, Tables 1 and S1). Deletion of the FAM32A C-terminus destabilized the 3′-ss and surrounding mRNA junction within the active site (Figure 4A-D). Focused classification revealed that progressive engagement of FAM32A residues 80-95 with the spliceosome correlates with Prp8 α-finger folding and partial docking of the 3′-ss, consistent with a model where the FAM32A tail stabilizes engagement of weaker, proximal 3′-ss in a late, catalytically competent docked configuration (Figure 4E).

**Figure 4.**
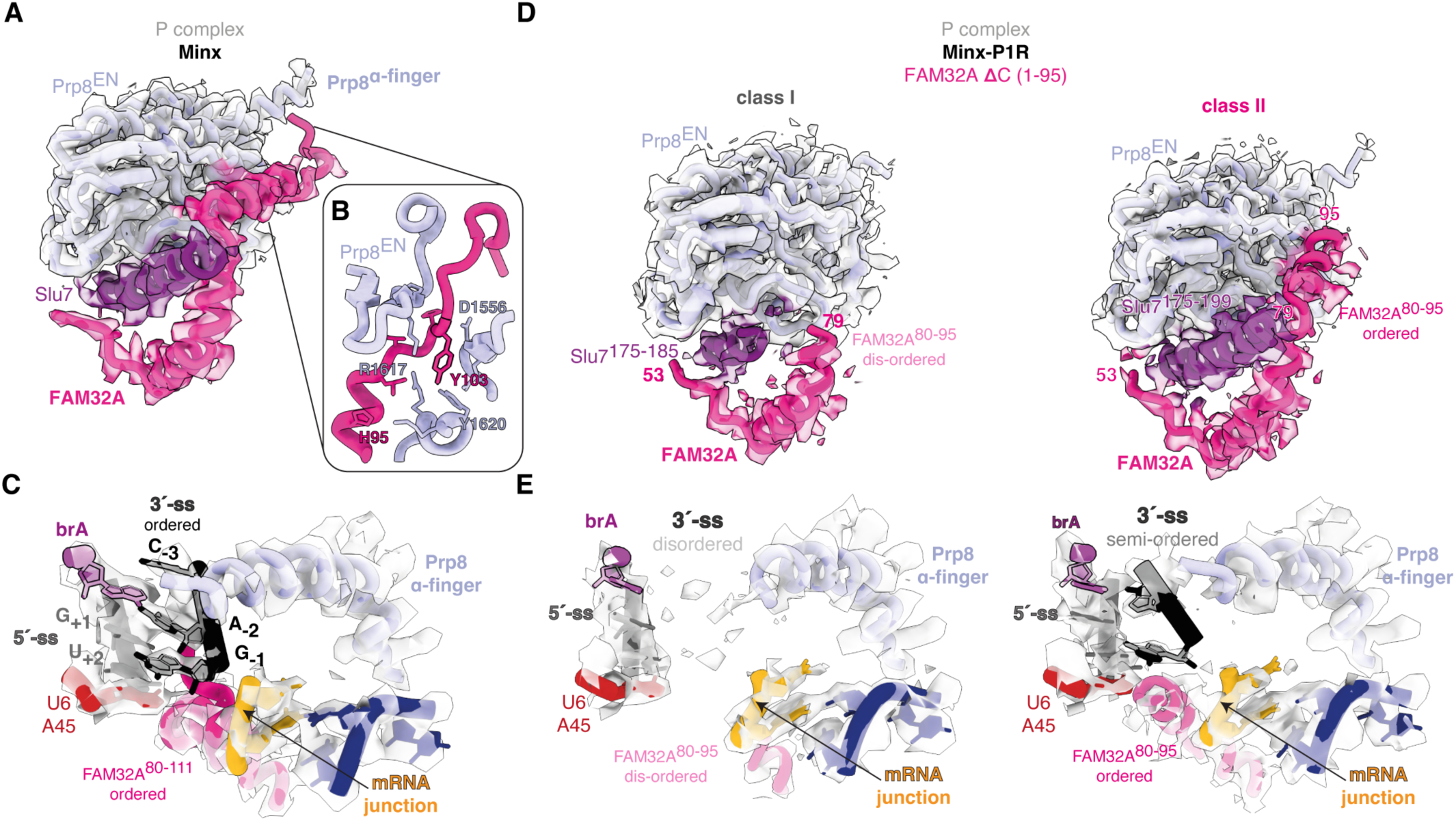
FAM32A stabilizes 3′-ss docking. **(A to C)** Overview of the human P-complex spliceosome on Minx pre-mRNA (PDB 6QDV) showing FAM32A and Slu7 bound alongside the Prp8 endonuclease (Prp8EN) and α-finger domains **(A)**, and anchoring of FAM32A **(B)**. Y103 serves as an anchor for the FAM32A C-terminus, guiding it into the active site. The FAM32A ΔC mutant is truncated at residue 95 and thus lacks this anchor, explaining why residues 80–95 are only ordered in a subset of particles. Density for the ordered 3′-ss and FAM32A C-terminus (80–111) in the Minx P complex is shown in **(C)**. **(D** and **E)** Progressive ordering of the FAM32A C-terminus correlates with increased ordering of Slu7 **(D)** and more stable docking of the 3′-ss **(E)**. Equivalent views for Minx-P1R P complexes show a class with a disordered 3′-ss lacking FAM32A 80–95 (class I) and a class with a semi-ordered 3′-ss showing density for FAM32A 80–95 (class II).

Together, these structural and functional data define a sequence-based code for 3′-ss selection, in which active-center geometry and the C* docking network act together to determine splice-site choice during catalysis.

### Perturbing C* docking factors corrects pathogenic mis-splicing *in vivo*

The *cis*-regulatory code for 3′-ss docking predicts that weak proximal sites will be highly dependent on exon-ligation factors for stabilization. Such arrangements of a weak proximal acceptor site followed by a stronger distal site are frequently observed for pathogenic mutations that introduce *de novo* AG dinucleotides upstream of canonical 3′-ss (AG-gain variants)^39^. At present, there are no splice-correcting strategies for such AG-gain variants, reflecting the previously limited molecular understanding of how competing 3′-ss are selected during catalysis.

To test whether our new molecular mechanism for 3′-ss selection can be applied to correcting AG-gain variants, we first used available literature and databases to identify pathogenic NAGNAG variants. Our *cis*-feature-based ML model predicted that 36 of the 43 identified variants would respond to perturbation of FAM32A (Figure 5A; Figure S10A). Using minigenes carrying pathogenic variants, we confirmed preferential disease-linked proximal selection in all tested AG-gain and PTC-creating NAGNAG sites (Figure 5A; Figures S10B-E). We selected predicted strong, intermediate and non-targets for investigating the response of these minigenes to exon-ligation factor perturbations. In high-confidence targets such as *SCN8A* disease-associated proximal site usage was almost fully reversed by FAM32A ΔC expression (Figure 5A; Figures S10B-E). Clinically relevant variants in *CFTR*, *OCRL*, and *BRCA1* were predicted to be less sensitive to FAM32A perturbation and indeed showed only modest rescue. Predicted non-targets like *SCN1A* remained completely insensitive to modulation of FAM32A and other exon-ligation factors, supporting both a sequence specific susceptibility to exon-ligation factor modulation and the predictive accuracy of our ML model (Figure S10B-E). Consistent with our sequence-based code for 3′-ss selection, the predicted responsiveness was strongly associated with a negative Δ splice-site score, reflecting a weak proximal and stronger distal splice site (Figure S10E).

**Figure 5.**
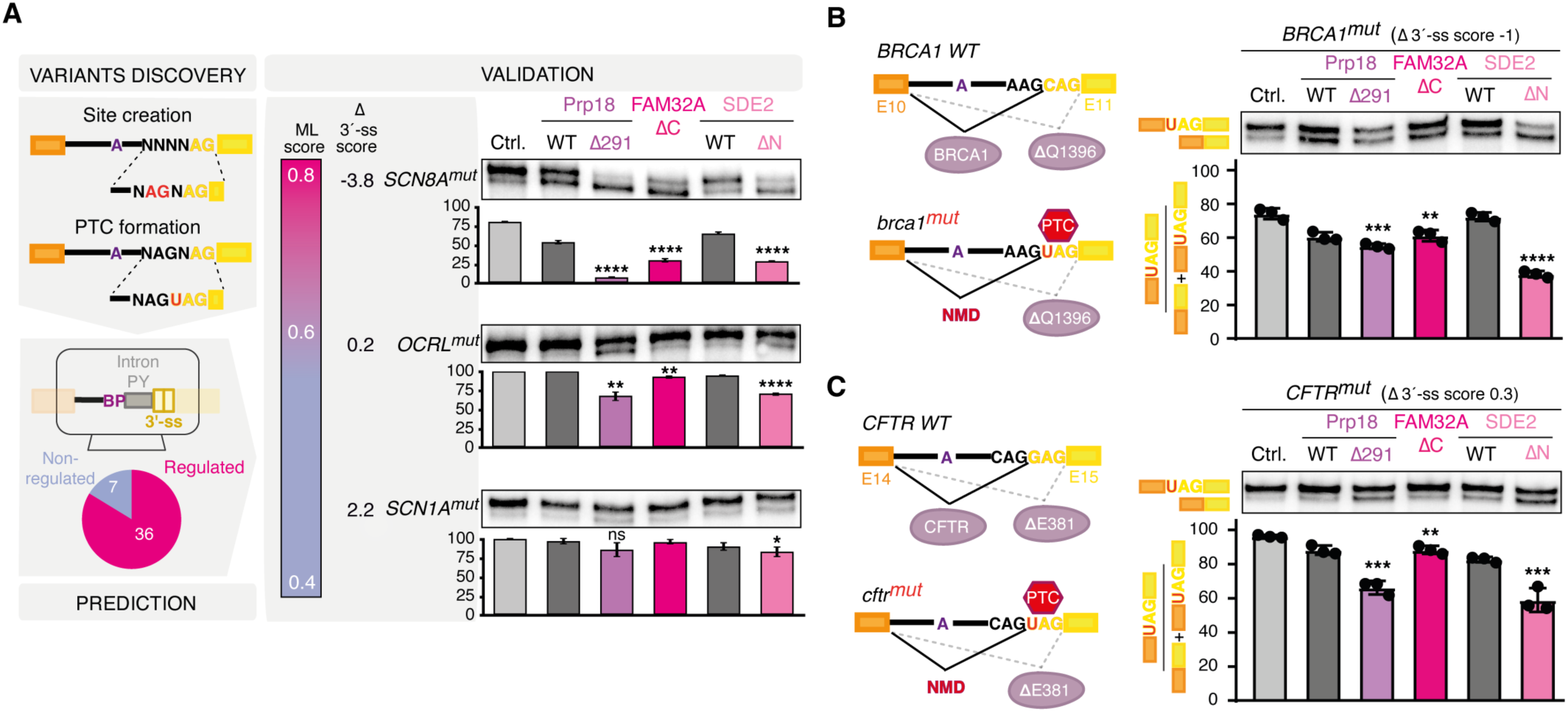
Perturbation of exon-ligation factors restores canonical splicing for pathogenic AG-gain variants. **(A)** Regulation of disease-related targets by exon-ligation factors can be accurately predicted by machine learning. Left: Selected single-nucleotide variants (SNVs) comprising AG-gain and PTC-creating mutations were predicted with our ML model; prediction rates are shown. Middle: Prediction score range and Δ 3′-ss scores for competing NAGNAG sites. Right: RT-PCR validation of select strong, intermediate or non-targets (*SCN8A*, *OCRL*, and *SCN1A*). **(B and C)** Pathogenic mis-splicing events can be differentially corrected by disrupting Prp18, FAM32A, or SDE2 function. Minigene assays are shown for a *BRCA1* 3′-ss mutation (*BRCA1* c.4186C>T) **(B)**, and a *CFTR* PTC-creating mutation in a cryptic 3′-ss (*CFTR* 2623G>T) **(C)**. Both mutations result in the formation of a new tandem 3′-ss that leads to a PTC-containing isoform subject to nonsense-mediated decay (NMD). Left schematics in each panel show wild-type and mutant 3′-ss structures. In all panels, data are represented as mean ± SD, with points indicating individual biological replicates (n = 3). Statistical significance is shown as indicated: * *p* < 0.05; *p* < 0.01; *** *p* < 0.001; **** *p* < 0.0001; ns, not significant.

Notably, clinically relevant NAGNAG variants introducing NMD-sensitive stop codons in *BRCA1* and *CFTR* remained highly responsive to other components of the docking network. Perturbation of Prp18 or SDE2 effectively suppressed the pathogenic proximal sites and restored canonical distal 3′-ss usage (Figures 5B, C; Figure S10E). Thus, distinct exon-ligation factors allow gene-specific correction of 3′-ss selection by the active center. Notably, despite being trained exclusively on FAM32A targets, our ML model predicts responsiveness to other components of the exon-ligation machinery, again pointing to a common mechanism, with the magnitude of the response being specific for individual exon-ligation factor perturbations. Together, these results demonstrate that disruption of the active-site docking network can reverse pathogenic 3′-ss selection for select AG-gain alleles, and establish 3′-ss docking during catalysis as a promising target for therapeutic intervention.

## Discussion

Our integrative study resolves the complete exon-ligation active site on a human intron derived from the native β-globin transcript and provides the structural and mechanistic framework for selection of competing 3′ splice sites during catalysis. We identify SDE2 as a previously unrecognized active-site constituent and show that, together with Prp18 and FAM32A, it stabilizes the exon-ligation conformation to promote efficient splicing and modulate 3′-ss choice.

Importantly, SDE2 stabilizes the branch helix for 3′-ss engagement following remodeling of the C complex (Figure 6A), allowing the BP and 5′-ss to pair with the docked 3′-ss (Figure S3F). Our cryo-EM structures of spliceosomes assembled with SDE2 ΔN directly visualize the scaffolding defect resulting from impairment of SDE2 function, revealing stalled C*-like complexes with a poorly ordered branch helix where other exon-ligation factors do not stably engage the spliceosome core (Figure 2C; Figure S3E). Intriguingly, because the SDE2 C-terminus stabilizes association of the disassembly ATPase DHX15 after mRNA release^40^, SDE2 may couple catalytic progression with kinetic surveillance, potentially targeting defective spliceosomes for disassembly (Figure 6A). Locally, SDE2 may be important for promoting K2 binding during 3′-ss selection, especially at weaker sites. Consistent with this, almost all tested pathogenic AG-gain variants respond to SDE2 N-terminus deletion (Figure S10E). These results uncover a fundamental splice site selection mechanism during splicing catalysis and provide the molecular explanation for SDE2 regulation of alternative splicing in humans^41^ and yeast^28^.

**Figure 6.**
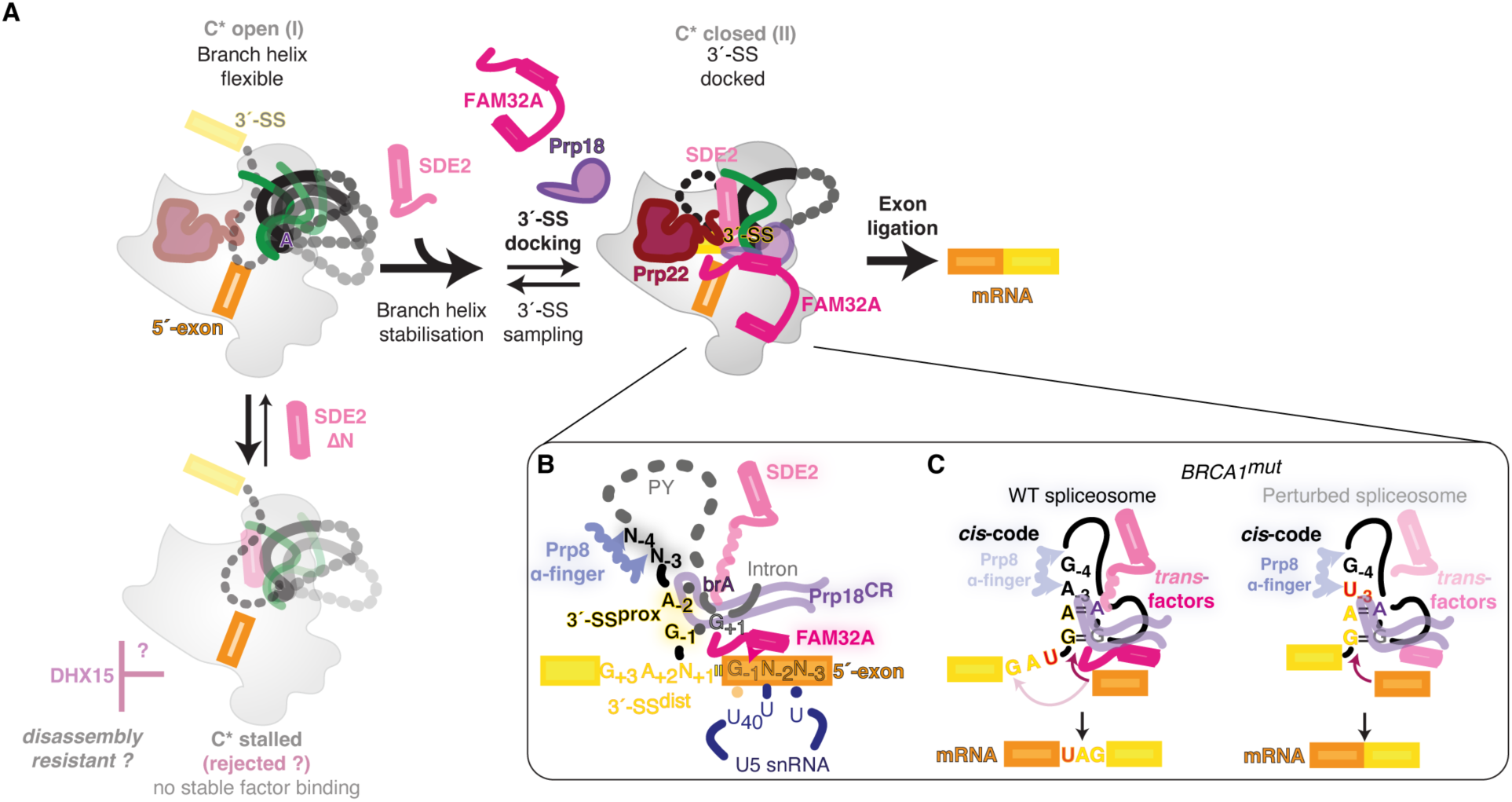
The spliceosome active center functions as a tunable regulatory hub for alternative 3′-ss selection. **(A)** Following C complex remodeling to C*, the SDE2 N-terminus rigidifies the branch helix, enabling transition to a docking-competent closed C* state. SDE2 ΔN stalls C* in an open state lacking stable factor binding. Loss of the SDE2 N-terminus is proposed to stall C* in this open conformation before stable 3′-ss docking, which may impair DHX15-mediated disassembly and promote transcript rejection (cf. ref. ^40^). **(B)** Active-site geometry. Exon-ligation factors cooperate with the Prp8 α-finger to stabilize the extended helical stack, modulating sampling between the proximal and distal 3′-ss. **(C)** *Trans*-acting factors modulate a *cis*-regulatory code. In wild-type spliceosomes, *trans*-factors dock the weaker, pathogenic proximal 3′-ss in *BRCA1* mutants against the intrinsic *cis*-code. Perturbing this *trans*-factor network destabilizes the proximal site, allowing the structural *cis*-bias to drive usage of the distal 3′-ss and produce canonical, non-pathogenic mRNAs.

By integrating cryo-EM, transcriptome profiling, and machine learning, we reveal that 3′-ss choice in the human C* spliceosome is governed by a sequence-based code that differentially stabilizes docking of competing 3′-ss (Figure 6B). This creates a tunable mechanism for regulation by exon-ligation factors, in which SDE2 promotes formation of a docking-competent C* complex. Within this conformation, contacts between the Prp8 α-finger and positions −4 and −3, together with the K2 site coordinating the docked 3′-ss, set the stability of the extended helix (Figure 6B). SDE2, FAM32A, and Prp18 then modulate the docked state, promoting stable engagement of weaker proximal 3′-ss that compete with stronger distal sites (Figure 6C). Interestingly, FAM32A also stabilizes Slu7 association with the spliceosome (Figure 4, D and E), potentially providing a mechanism to explain the reported role of Slu7 in alternative 3′-ss selection^42^. Thus, C* docking factors act cooperatively with other exon-ligation factors to ensure accurate 3′-ss selection and efficient catalytic progression.

This mechanism suggests that exon-ligation factors act to stabilize the C* spliceosome at the first AG downstream of the branch point and constrain downstream competition, in agreement with previous models of 3′-ss scanning^43^. Within this framework, Prp22 ATPase activity may further promote sampling of alternative 3′-ss, as observed previously in yeast^44^, raising the question of how its proofreading role adapts to human splicing complexity. The existence of *trans*-factors like SDE2 and FAM32A that act as “readers” of a *cis*-code correlates with splicing complexity^45^, but their regulatory mechanism creates vulnerability to pathogenic AG-creating mutations. However, by using perturbation of exon-ligation factors to restore canonical splicing in disease-relevant contexts, we establish 3′-ss docking as a tractable, structure-guided target for therapeutic interventions to correct pathogenic mis-splicing in diverse AG-gain mutants.

## Acknowledgements

We thank S. Scheres for his help and advice on data collection and processing; S. Chen, G. Cannone, J. Grimmett, and T. Darling for smooth running of the EM and computing facilities at the LMB; R. Matadeen and E. Lowe for help with data collection and processing at COSMIC; the staff at Diamond Light Source (DLS) for help with data collection; the MRC-LMB and Oxford Biochemistry mass spectrometry facilities for help with protein identification; S. Yang for purification of Strep-tagged MS2-MBP protein; N. Proudfoot, G. Dujardin, and R. DeSousa-Louis for advice and support. V.-Y. Feng and S. R. Reyna for help with minigene experiments; and the HPC Service of FUB-IT, Freie Universität Berlin, for computing time. We thank W. Galej, N. Proudfoot, R. Klose, and F. Mardakheh for critical reading of the manuscript and C. Smith for a generous gift of reagents.

The project was supported by the Medical Research Council (MC_U105184330), a European Research Council Advanced Grant (SPLICE3D), and the Wellcome Trust (ALR02270).

S.M.F. was supported by a Marie Skłodowska-Curie fellowship, and by a Wellcome Trust and Royal Society Sir Henry Dale Fellowship (ALR02270).

F.H. is supported by the Deutsche Forschungsgemeinschaft (DFG, German Research Foundation) under Germany’s Excellence Strategy – EXC 3118/1 – project number 533770413.

## Author contributions

S.M.F. initiated the cryo-EM project, discovered SDE2 binding in the spliceosome, analyzed the structural data, and coordinated and supervised cryo-EM and biochemistry experiments. M.P. and F.H. initiated, coordinated, and supervised the functional experiments *in vivo*. S.M.F. established the method of P complex preparation, designed the substrates used for cryo-EM studies, prepared large-scale nuclear extracts, purified Prp22 mutants for cryo-EM, prepared EM samples and grids for the β-globin and Minx-P1R samples, collected and performed initial processing of cryo-EM data, discovered the new C* conformation in the SDE2 ΔN dataset, and performed most *in vitro* splicing assays. G.M. purified SDE2, Prp18, and FAM32A, re-processed the full cryo-EM datasets, carried out model building, and refined and deposited the maps and PDB files. T.M. prepared nuclear extracts, and purified the samples and collected the cryo-EM datasets for the SDE2 ΔN complexes. L.Y. prepared splicing substrates and nuclear extracts, purified Prp18 and mutant Prp22, and performed *in vitro* assembly assays, glycerol gradients, and SDE2 EMSA experiments. Y.S. prepared graphene grids for cryo-EM and helped with grid screening. L.C. performed SDE2 EMSA experiments. G.-V. D. and L.Y. performed small scale spliceosome assembly and Western blotting for Prp18. S.E. performed all minigene and FACS experiments with help from M.P. and H.Y.K. L.Z. and M.P. analyzed RNA-seq data. L.Z. created and trained the machine learning model, and identified pathogenic variants. S.M.F., M.P., and F.H. wrote the manuscript with input from all authors.

## Data and code availability

The cryo-EM maps have been deposited in the Electron Microscopy Data Bank, and the coordinates of the atomic models have been deposited in the Protein Data Bank. Accession codes will be made available upon peer-reviewed publication of the manuscript. RNA-seq data are available at the European Nucleotide Archive under PRJEB101467. Code and required data for RNA sequencing and machine learning analyses are available at GitHub (https://github.com/luizazuvanov/human-a3ss-control-pathogenic-missplicing).

## Methods

### Plasmid construction

To generate pSMF71 (encoding hFAM32A (2–95) with an N-terminal hexahistidine [His_6_]–glutathione S-transferase [GST]–TEV cleavage site and a Strep II tag), human *FAM32A* (UniProt accession no. Q9Y421) was amplified from HeLa cell cDNA (Takara) using primers oSMF275/oSMF277 and cloned into the pGEX-4T-1 vector linearized with primers oSMF273/oSMF274 using the NEBuilder HiFi DNA Assembly Cloning Kit (New England Biolabs), following the manufacturer’s instructions.

To generate pSMF86 (encoding hFAM32A^2–112^ with an N-terminal TEV-cleavable His_6_–GST tag, followed by a Strep II tag after the TEV site), *FAM32A* was amplified from pSMF71 using primers oSMF390/oSMF391 and cloned into pGEX-4T-1 as above.

To generate pSMF103 (encoding hSDE2^78–451^ with an N-terminal SUMO tag and a C-terminal TEV-cleavable Strep II tag), *SDE2* (UniProt accession no. Q6IQ49) was amplified from HeLa cell cDNA using primers oSMF490/ oSMF491, and cloned into a pET-Duet vector linearized with primers oSMF435/oSMF492 using the NEBuilder HiFi DNA Assembly Kit.

To generate pSMF107 (encoding hSDE2^95–451^ with an N-terminal SUMO tag and a C-terminal TEV-cleavable Strep II tag), the SUMO tag was amplified from pSMF103 using primers oSMF490/oSMF506 and the *SDE2* fragment with primers oSMF505/oSMF491. The two fragments were assembled into pET-Duet linearized with oSMF435/oSMF492 using NEBuilder HiFi DNA Assembly.

The coding sequence for human Prp18 was amplified from HeLa cDNA (Sigma) and cloned into a pGEX-4T-1 vector containing an N-terminal His-GST-TEV-Strep II tag.

The coding sequence for human Prp22 (DHX8) was amplified from HeLa cDNA (Sigma) and cloned into a vector containing an N-terminal FLAG-TEV-Strep II tag (pWG219, a gift from W. Galej), to generate pSMF8. The K594A and the C-terminal deletion mutants were introduced by site-directed mutagenesis as described^46^.

An MS2-MBP variant with an additional N-terminal twin Strep-tactin tag was a gift from Chris Smith (Cambridge University).

For expression in human cells, codon-optimized cDNA sequences of FAM32A (ref. ^25)^ and Prp18-each bearing a N-terminal FLAG tag - as well as SDE2 were ordered in the pTWIST-CMV expression vector (Twist Bioscience). PCR products were integrated into appropriate vectors using restriction enzymes.

To generate the FAM32A ΔC (1–H95) construct, the truncated fragment was amplified using primers PC-001 and PC-002 and cloned into the pTWIST-CMV vector.

For generation of the Prp18 ΔR291 construct, two fragments were amplified using primer pairs PC-032/PC-035 and PC-034/PC-033. The fragments were fused by overlap extension PCR and subsequently amplified using primers PC-032 and PC-033. The resulting product containing the deletion was cloned into the pTWIST-CMV vector.

To obtain SDE2 WT with a C-terminal FLAG tag, the tag sequence was introduced using primers PC-085 and PC-090. SDE2 variants were generated using primers PC-087/PC-090 to produce the SDE2 ΔN (90–K451) construct and primers PC-086/PC-090 to generate the SDE2 Met-K78 (78–K451) construct. All variants were cloned into the pTWIST-CMV expression vector.

The sequence for b-globin pre-mRNA was cloned into a pUC19 vector backbone from RR108 (a gift from C. Smith ^47^); for spliceosome assembly the pre-mRNA flanking exons were truncated to 45 nucleotides to match the Minx pre-mRNA exon length. The Minx-P1R chimera was constructed by cloning the polypyrimidine tract of a *PPP1R12C* intron^25^ into the standard Minx construct described previously^48^.

For minigene experiments, the BP-to-3′ss sequences of endogenous or disease-associated introns were introduced by PCR into a GFP interrupted by an intron ^49^. In wildtype or mutated AG-gain variants, the last 50 nt of the 3′-ss together with first 10 nt of the downstream exon were cloned into RG6 open reading frame ordered in pTWIST. Additionally, the entire *PPP1R12C* NAGNAG intron sequence, with surrounding exons, was amplified from genomic DNA and cloned into pcDNA3.1+. For FACS experiments, the *PPP1R12C* 3′ splice site sequence was inserted into the GFP intron, and the distal CAG was mutated to generate an in-frame stop codon (UAG).

### Preparation of Splicing Substrates

Pre-mRNA for C*/P complex assembly have three copies of MS2 coat protein aptamers cloned at the 3′-end, whereas substrates for biochemical studies do not. Substrate mutations were introduced by site-directed mutagenesis as described^46^. Transcription templates were prepared by PCR from the pre-mRNA plasmids and transcribed with in-house purified T7 RNA polymerase. The transcript was purified by ion exchange chromatography using a HiTrap DEAE Sepharose FF column (Cytiva) followed by a NAP-10 column (GE Healthcare) for buffer exchange. The purified transcript was capped using the Vaccinia capping system (NEB) and 3′-end labelled using T4 RNA Ligase 1 and pCp-Cy5 (Jena Bioscience) according to the manufacturer’s instructions, with each step followed by buffer exchange using G25 columns (GE Healthcare).

### HeLa Nuclear Extract Preparation and in vitro Splicing

8-16 L of HeLa cells were grown to a density of 0.6 – 1 million cells / mL at 37 °C with 5% CO_2_ and agitated at 120 rpm. Nuclear extract was prepared using a 30 mL Dounce homogenizer in batches essentially as described^50^, and dialyzed against buffer D100 (20 mM HEPES, pH 7.9, 100 mM KCl, 20% glycerol, 0.2 mM EDTA).

*In vitro* splicing was performed essentially as described^51^, with 40% nuclear extract in the presence of 2 mM ATP, 20 mM Creatine Phosphate (Roche), and 3mM MgCl_2_. For C*/P Complex assembly, 12 nM pre-mRNA was used for the reaction. For functional studies, 24 nM pre-mRNA was used. Prp22 was added to a final concentration of ∼ 0.2 μM, Prp18 was added to a final concentration of ∼ 0.9 μM, FAM32A was added at a concentration of ∼ 1.2 uM, and SDE2 was added at ∼ 45-90 nM in the splicing reaction.

### Purification of Recombinant Proteins

FAM32A, SDE2, and Prp18 plasmids were transformed into *Rosetta (DE3) pLysS* cells and expressed in 2 L of Power Broth media (Molecular Dimension). FAM32A constructs were induced at OD_600_ ∼0.6 with IPTG and grown for 5 h at 30 °C. SDE2 constructs were induced overnight at OD_600_ ∼0.7 at 28 °C. Prp18 constructs were induced at OD_600_ ∼0.6 with IPTG and grown overnight at 30 °C.

#### FAM32A purification

Cells were harvested by centrifugation and resuspended in 25 mL lysis buffer per liter of culture (50 mM HEPES, pH 7.5, 500 mM NaCl, 1 mM DTT, and one cOmplete™ Protease Inhibitor Tablet per 60 mL). Cells were lysed by sonication, and lysates were clarified by centrifugation at 45,000 × *g* for 30 min at 4 °C. The supernatant was incubated with 1.5 mL GST resin (Cytiva), washed with 15 mL lysis buffer, and the protein was eluted by overnight on-column cleavage with in-house purified TEV protease in 1.5 mL lysis buffer at 4 °C. The TEV-cleaved protein was applied to 0.5 mL Strep-Tactin® XT 4Flow® resin (IBA Bioscience), washed sequentially with 3 mL lysis buffer and 3 mL low-salt buffer (20 mM HEPES, pH 7.9, 200 mM KCl, 1 mM DTT), and eluted with 4 mL Strep elution buffer (20 mM HEPES, pH 7.9, 200 mM KCl, 60 mM biotin, 1 mM DTT). The salt concentration was reduced to 50 mM NaCl using buffer A (50 mM HEPES, pH 7.9, 1 mM DTT), and the sample was loaded onto a 1 mL HiTrap SP XL column (Cytiva) equilibrated in 5% buffer B (50 mM HEPES, pH 7.9, 1 M NaCl, 1 mM DTT). Protein was eluted with a gradient from 50 mM to 1 M NaCl. Fractions were pooled, dialyzed into buffer D (20 mM HEPES, pH 7.9, 200 mM KCl, 20% glycerol, 1 mM DTT), and stored at –80 °C. Protein concentration was determined by A_280_.

#### SDE2 purification

Cells were harvested and resuspended in 25 mL lysis buffer per liter of culture (20 mM HEPES, pH 7.9, 500 mM NaCl, 2 mM DTT, and one cOmplete™ Protease Inhibitor Tablet per 60 mL). Cells were lysed by sonication, and lysates were clarified by centrifugation at 45,000 × *g* for 30 min at 4 °C. The supernatant was incubated with 0.2 mL Strep-Tactin® XT 4Flow® resin (IBA Bioscience), washed with 3 mL lysis buffer and 3 mL low-salt buffer (20 mM HEPES, pH 7.9, 200 mM KCl, 2 mM DTT). The SUMO tag was removed by on-column cleavage with SENP1 protease (1:10 v/v of a 3 mg/mL stock) overnight at 4 °C in 0.2 mL low-salt buffer. SENP1 cleaves after *GG* in the *GG*|KGG, thus leaving a clean KGG N-terminus for SDE2; the resulting protein mass was confirmed by mass spectrometry (data not shown). Beads were washed with 0.6 mL low-salt buffer and eluted with 1 mL Strep elution buffer (20 mM HEPES, pH 7.9, 200 mM KCl, 60 mM biotin, 2 mM DTT). Eluted protein was subjected to size-exclusion chromatography on a Superdex 200 Increase 10/300 GL column (Cytiva) equilibrated in 20 mM HEPES, pH 7.9, 200 mM KCl, 2 mM DTT. Peak fractions were pooled, dialyzed into buffer D (20 mM HEPES, pH 7.9, 200 mM KCl, 20% glycerol, 1 mM DTT), and stored at –80 °C. Protein concentration was determined by A_280_.

#### Prp18 purification

Cells were harvested by centrifugation and resuspended in 25 mL lysis buffer per litre of culture (50 mM HEPES, pH 7.5, 500 mM NaCl, 1 mM DTT, and one Complete™ Protease Inhibitor Tablet per 60 mL). Cells were lysed by sonication, and lysates were clarified by centrifugation at 45,000 g for 30 mins at 4 °C. The supernatant was incubated with 1 mL Glutathione Sepharose beads (Cytiva), washed with 10 mL lysis buffer, and the protein was eluted by overnight on-column cleavage with 1:50 v/v TEV protease in 1 mL lysis buffer at 4 °C. The TEV-cleaved protein was applied to 0.4 mL Strep-Tactin® XT 4Flow® resin (IBA Bioscience), washed with 3 mL lysis buffer and 3 mL wash buffer (20 mM HEPES, pH 7.9, 200 mM KCl, 1 mM DTT), and eluted with 4 mL Strep elution buffer (20 mM HEPES, pH 7.9, 200 mM KCl, 1 mM DTT, 10% Glycerol, 60 mM biotin). The protein was dialyzed into buffer D (20 mM HEPES, pH 7.9, 200 mM KCl, 20% glycerol, 1 mM DTT) and stored at −70 °C.

#### Prp22 purification

The wild-type and mutant Prp22 (DHX8) were expressed in HEK293 cells. Cells were harvested by centrifugation and resuspended in 25 mL lysis buffer (50 mM HEPES, pH 7.5, 500 mM NaCl, 1 mM DTT, and one cOmplete™ Protease Inhibitor Tablet per 60 mL) per 400 mL of culture, then lysed by sonication. The lysates were clarified by centrifugation at 45,000 g for 30 mins at 4 °C and the supernatant was incubated with 0.2 mL Strep-Tactin® XT 4Flow® resin (IBA Bioscience), washed sequentially with 15 mL NaCl wash buffer (20 mM HEPES, pH 7.9, 500 mM NaCl, 10 mM β-Mercaptoethanol, 10% Glycerol) and 8 mL KCl wash buffer (20 mM HEPES, pH 7.9, 200 mM KCl, 10 mM β-Mercaptoethanol, 10% Glycerol), and eluted with 3 mL Strep Elution buffer (20 mM HEPES, pH 7.9, 200 mM KCl, 10 mM β-Mercaptoethanol, 10% Glycerol, 60 mM biotin). The purified proteins were dialyzed into buffer D (20 mM HEPES, pH 7.9, 200 mM KCl, 20% glycerol, 1 mM DTT) and stored at −70 °C.

### Analytical C*/P complex assembly

Spliceosome complexes were assembled on MINX and β-globin pre-mRNA substrates pre-bound to MS2-MBP fusion protein in 1.25 - 2.5 mL *in vitro* splicing reactions supplemented with recombinant hPrp22 (DHX8) K594A. Reactions were incubated at 30 °C for 60 minutes (MINX) or 90 minutes (β-globin). DHX8 K594A mutant prevents the release of mRNA and stalls the spliceosome at C*/P complex, thereby protecting the 3′-exon from RNase H-mediated cleavage. To remove spliceosomes without a docked 3′-exon, the splicing reaction was supplemented with 4 μM of DNA oligonucleotide complementary to the 3′-exon and incubated for an additional 10 minutes to induce cleavage of the 3′-MS2 tag by the endogenous RNaseH activity of the splicing extract. Protected C*/P complexes were purified through MS2-MBP fusion protein, using buffers containing 100 mM or 150 mM KCl and 0.025% NP40 substitute. Fractions containing spliceosomes were determined by Cy5 fluorescence, pooled, and concentrated by progressive re-spinning in 250 μL aliquots in a TLA100 rotor at 42000 rpm to reduce the volume. Spliceosomes migrated into the bottom 50-75 μL after one hour.

### Western Blotting

For Western Blotting from *in vitro* complex assemblies, primary antibodies against Snu114 (GeneTex), Prp18 (ABclonal), and Slu7 (Santa Cruz Biotechnology) were used at dilutions of 1:4000 (Snu114 and Prp18) or 1:20,000 (Slu7). Detection was carried out with HRP-conjugated secondary antibodies (goat anti-rabbit, Invitrogen, 1:5000; goat anti-mouse, Promega, 1:5000) and visualized using enhanced chemiluminescence (ECL).

For Western blot analysis of overexpressed C* proteins, approximately 3.75 × 10⁵ HEK293 cells were seeded in six-well plates. Twenty-four hours after seeding, cells were transfected with 2 µg plasmid DNA encoding FLAG-tagged proteins using 5 µl ROTI®Fect (Carl Roth).

Forty-eight hours post-transfection, cells were washed with 1× PBS and lysed in 1× RIPA buffer (10 mM Tris-HCl pH 7.5, 100 mM NaCl, 2 mM EDTA, 1% NP-40) supplemented with protease inhibitors. Cell debris was removed by centrifugation, and the supernatant was boiled in SDS-sample buffer.

Whole-cell lysates were separated by 12 % SDS-PAGE and transferred to nitrocellulose membranes via semi-dry blotting. After blocking in 2 % BSA, dissolved in 1x TBST, membranes were incubated with rabbit polyclonal anti-FLAG antibody (DYKDDDDK tag, Cell Signaling Technology) at 1:4000 dilution and mouse monoclonal anti-HNRNPL antibody (clone 4D11, Santa Cruz Biotechnology) at 1:10,000 dilution. HRP-conjugated secondary antibodies were applied at 1:2000 dilution and signals were detected by enhanced chemiluminescence (ECL).

### Splicing factor depletion and reconstitution and analytical Western Blotting

FAM32A was depleted from HeLa S3 nuclear extracts prepared fresh, essentially as described 48, using affinity-purified antibodies against FAM32A Abcam (Abcam 185298). Depletion was at least 90%, as judged by Western Blotting using the same anti-FAM32A antibody as for depletion (Figure S5E). For reconstitution of activity in FAM32A-depleted extract, recombinant FAM32A was added at a concentration of 0.6-1.2 uM.

Prp18 was depleted from HeLa S3 nuclear extracts prepared fresh, essentially as described35, except that Prp18 antibodies (Abcam 12817) were crosslinked to Protein-A-MagSepharose Xtra (GE Healthcare) using DMP prior to immunodepletion. For double immunodepletions, Prp18 was depleted first following by depletion of FAM32A. Depletion was at least 90%, as judged by Western Blotting using the same Prp18 antibody as for depletion (Figure S5, G and I).

SDE2 was depleted from HeLa S3 nuclear extracts prepared fresh, using SDE2 antibodies (Bethyl Laboratories A302-098A) pre-conjugated to Protein-A-MagSepharose Xtra (GE Healthcare), and crosslinked with DMP prior to depletion. Approximately 25 uL of antibody (1 mg/mL) was used for every 65 uL of splicing extract. Depletion was 50-60%, as judged by Western Blotting using the same SDE2 antibody as for depletion (Figure S3G).

For Western blotting, primary antibodies (FAM32A from Abcam (185298); Prp8 from Proteintech; Slu7 from SantaCruz Biotech (D3)) were used at dilutions of 1:200-1:1000 and detected by ECL-conjugated Donkey-anti-Rabbit (FAM32A and Prp8) at 1:1000 or anti-mouse (GE) at 1:5000 secondary antibodies using the ECL system (GE Healthcare).

### Cell culture, transfection and siRNA knockdown

HEK293T cells (ATCC® CRL-1573™) were cultured in Dulbecco’s Modified Eagle Medium (DMEM) supplemented with 10% fetal bovine serum and 100 µg/ml penicillin–streptomycin. Mycoplasma contamination was routinely monitored by monthly PCR-based testing and assessment of cell morphology.

For plasmid transfection, approximately 1.5 × 10⁵ cells per well were seeded into 12-well plates 24 h prior to transfection. Transfections were performed using ROTI®Fect (Carl Roth) with 4 µl reagent per well for RNA-seq experiments and 2 µl per well for minigene assays and FACS analysis.

For RNA-seq experiments, 1.2 µg plasmid DNA was transfected per well. For minigene co-expression experiments, 0.2 µg minigene plasmid and 0.6 µg protein expression plasmid were co-transfected. Cells were harvested 48 h post-transfection.

For FACS experiments, 0.6 µg SDE2 WT, ΔN or Met-K78 expression constructs in pTWIST-CMV were co-transfected with 0.2 µg GFP-CAG or GFP-CAGUAG minigene plasmids per well. Cells were harvested 24 h after transfection.

For siRNA knockdown experiments, 5 × 10⁴ HEK293T cells were seeded per well in a 12-well plate and transfected with 20 pmol siRNA targeting U2AF35 (GAAAGUGUUGUAGUUGAUUGA) or FAM32A (GGAGAAGCGGCAAAUGGAA), or a control siRNA (UUCUCCGAACGUGUCACGU). Cells were harvested 72 h post-transfection and total RNA was extracted.

### RNA extraction, RT-PCR and RT-pPCR

To analyze NAGNAG alternative splicing events, RNA extraction and RT-PCR were performed as described previously^52^. HEK293 cells were harvested 48 h post-transfection using RNATri reagent (Bio&Sell), and total RNA was isolated. Residual genomic DNA was removed by DNase I digestion. RNA quality and concentration was measured using nanodrop photometer.

For reverse transcription, 1 µg total RNA was subjected to gene- or minigene-specific RT using up to four reverse primers targeting different transcripts in a single reaction. Forward primers used in subsequent PCR reactions were labelled with [γ-³²P]-ATP to enable sensitive detection of splice isoforms.

PCR products were separated on 6% denaturing polyacrylamide gels. To achieve optimal resolution of NAGNAG splice isoforms, PCR primers were designed to generate amplicons of 90–150 nt. Bands were visualized using a PhosphorImager and quantified with ImageQuant TL (Cytiva).

Quantified values represent mean ± standard deviation. Statistical significance was assessed using an unpaired Student’s t-test (*, p < 0.05: **, p < 0.01; ***, p < 0.001; ****, p<0.0001).

For RT-qPCR analysis of knockdown efficiency, gene-specific primers for *U2AF35* and *FAM32A* were used together with a housekeeping gene primer in the RT reaction. Quantitative PCR was performed using the Blue S’Green qPCR Kit Separate ROX (Biozym) on a Stratagene Mx3000P instrument. Reactions were performed in technical duplicates and normalized to hGAPDH expression.

### Flow cytometry

Cells expressing the GFP reporter were harvested using trypsin and washed once with 1× PBS. GFP fluorescence intensity was measured using a Guava easyCyte 8 flow cytometer at 488 nm. Untransfected cells served as a negative control.

For each sample, 20,000 events were acquired. Data were analyzed using GuavaSoft (version 2.7). A gating strategy excluding debris and doublets was applied.

Median GFP fluorescence intensity of cells expressing SDE2 mutant constructs was normalized to cells expressing wild-type SDE2 within each experiment. GFP values are presented as normalized fluorescence intensities. Error bars represent the standard deviation of independent experiments. Statistical significance was calculated using an unpaired Student’s t-test (*, p < 0.05; **, p < 0.01; ***, p < 0.001; ****, p < 0.0001).

### RNA-seq and data analysis

RNA-seq was performed in biological triplicates using DNase I–digested RNA samples for library preparation. Libraries were prepared using poly(A)+ selection at BGI Genomics and sequenced using DNBSeq PE150 sequencing. Each overexpression sample was compared to a CTRL transfection triplicate that was prepared and sequenced on the same day. C* factor mutant overexpression samples were additionally compared to WT variant over-expression. Read trimming and quality control were done using fastp (version 0.23.4, ref.^53^). Cleaned reads were aligned to the human genome (version GRCh38) using STAR (version 2.7.9, ref.^54^). Alternative splicing analysis was done with rMATS-turbo (version 4.1.1, ref.^55^). Events were considered significant if supported by a mean read coverage ≥ 10 and a False Discovery Rate (FDR) ≤ 0.01. To increase the confidence of calls more susceptible to background noise, mapping ambiguity, and unprocessed transcript contamination^56^, an additional filter with an absolute Inclusion Level Difference ≥ 0.15 was applied to intron retention events.

### Machine learning model

Data selection. rMATS alternative splicing analysis from RNA-seq data of siFAM32A and respective control samples, as previously described^25^, was used to classify NAGNAG events as being regulated by FAM32A or not. We used siRNA-mediated knockdown rather than mutant overexpression, as the 72 h knockdown samples were sequenced at greater depth, resulting in more pronounced effects on endogenous NAGNAG splice site choice. For highly confident events, NAGNAG was considered regulated if FDR ≤ 0.01, abs(dPSI) ≥ 0.15, and mean coverage reads ≥ 10 (statistics from rMATS software). NAGNAG was classified as non-regulated if mean coverage reads ≥ 10, abs(dPSI) ≤ 0.05, and TOST-FDR ≤ 0.01 (statistics were computed using a TOST-equivalent test, with p-values adjusted by the Benjamini-Hochberg method for False Discovery Rate). Information from the upstream exon, intron, downstream exon, inclusion level and gene expression was retrieved from the two NAGNAG groups using a Python script. Branch point information and 3′-ss scores were obtained with SVM-BPfinder^57^ and MaxentScan ^58^ software, respectively. Gene expression information was obtained using gene counts calculated previously during STAR alignment (version 2.7.9)^54^ followed by GeTMM normalisation with edgeR (version 4.6.3) in a homemade R script (version 4.5.1) to enable both within- and between-sample comparisons ^59^. See the “Data and Code Availability” section for further details about the code and input data sets used for data selection.

#### Model architecture

Sequence-retrieved features from the regulated and non-regulated NAGNAG events were used for machine learning. Three models were trained with different feature sets: (i) all sequence-retrieved features, (ii) “3’-splice site” together with “branch point” related features, or (iii) “5’-splice site” and “local context” related features. Using each set of features, a binary logistic base XGBoost model^60^ was trained on 70% of a stratified random split of the input dataset. Logarithmic loss (log loss) was used as XGBoost’s internal evaluation metric. Hyperparameter tuning of the base model was performed using Optuna’s Tree-structured Parzen Estimator (TPE) algorithm integrated with repeated stratified cross-validation (5 folds, 5 repeats)^61^. Matthews correlation coefficient (MCC) was used as the optimization objective and maximized across each of Optuna’s trials. Fifty optimization trials were performed within a hyperparameter search space including the number of trees (50-200), maximum tree depth (3-10), learning rate (0.01-0.3, log-scaled), subsampling ratio (0.5-1.0), column subsampling ratio (0.5-1.0), minimum loss reduction (gamma, 0-5), and minimum child weight (1-10). A final XGBoost model was retrained on the full training dataset using the optimal hyperparameters. The optimized model performance was assessed using repeated stratified cross-validation on the training set. In addition, to interpret model behavior and quantify feature contributions, SHapley Additive exPlanations (SHAP) values were computed for the optimized model using the training dataset^62^. The optimized model was later evaluated on an independent test set comprising the remaining 30% of the input data withheld from training. Performance metrics included MCC, accuracy, precision, recall, F1-score, and log loss. See the “Data and Code Availability” section for further details about requirements, code, input and output datasets files from model training, hyperparameter optimization, evaluation, and SHAP-based interpretability analyses.

### Annotated TASS variants

Selected variants comprise either mutations within (i) existing NAGNAG sites or mutations leading to the formation of (ii) novel NAGNAG sites. (i) Disease-related single-nucleotide variants (SNVs) introducing a premature stop codon (PTC) within existing distal NAGNAG splice sites were retrieved from the ClinVar database (downloaded on July 17th, 2025 ref. ^63^) and available literature^14^. (ii) Variants creating novel NAGNAG sites, defined by intronic variants with an introduced AG dinucleotide between the branch point and the 3’ splice site, were obtained from AGAIN’s publicly available dataset^15^. The selected dataset includes experimentally verified variants whose genes were clinically associated with diseases, and variants originate from genes implicated in cancer initiation and progression. A not directly disease-associated variant, located in the *SCN8A* gene, and a not yet proven pathogenic variant located in a critical region of the gene *IRF7* (both derived from AGAIN’s database) were also included in our pool. Sequences from variants (i) and (ii) were cloned into a minigene system as previously described and used for predicting their response to impaired FAM32A function using our machine learning model.

### Electron cryo-microscopy

#### C* / P complex assembly and EM sample preparation

Complexes were assembled on the b-globin or MINX-P1R pre-mRNA substrate pre-bound to MS2-MBP, or StrepII-MS2-MBP, fusion protein, respectively, in 50-120 mL *in vitro* splicing reactions supplemented with 0.03 mg/ mL of dominant-negative hPrp22 (DHX8) K594A mutant protein. For Minx-P1R, FAM32A lacking the C-terminal 17 amino-acids (2-95) at a final concentration of ∼ 1.2 uM was also added. Reactions were incubated for 90 minutes (for b-globin) or 60-90 minutes (for Minx-P1R chimeras) at 30°C. DHX8 K594A stalls the spliceosome at the C*/P complex stage by blocking release of the mRNA, which is protected from RNaseH cleavage directed against the 3′-exon. To remove spliceosomes without a docked 3′-exon, the splicing reaction was incubated for an additional 10-12 minutes with 4-5 mM of DNA oligonucleotide complementary to the 3’-exon: for Minx RH3 5’-CGTCCTCAACCGCGAG-3 ‘; for b-globin 5 ‘-GTAGACCACCAGCAGC-3’; for Minx-P 1 R RH 5 5 ‘CCTCAACCGCGAGCTG-3’. This procedures induces cleavage of the 3’-MS2 tag by the endogenous RNaseH activity of the splicing extract. Protected C*/P complexes were then purified via MS2-MBP fusion protein, essentially as described^48^, using buffer containing 100 mM KCl and 0.025% NP-40 substitute. For the C* / P complex assembled with DHX8 K594A on Minx-P1R pre-mRNA, a second affinity step using the MS2-MBP Strep-tactin tag was performed from the pooled maltose elution fractions, and enriched complexes were eluted with 5 mM DSB. Fractions containing lariat-intermediates and mRNA, as determined by Cy5 fluorescence, were pooled and concentrated by progressive re-spinning in 250 uL aliquots in a TLA100 rotor at 42000 rpm to reduce the volume, as described previously^48^. Samples were dialysed against buffer K100 (20 mM HEPES, pH 7.9, 100 mM KCl, 0.2 mM EDTA) 4 times for 45 minutes each in cassettes with a 20 kDa MWCO. For spliceosomes assembled in the presence of SDE2 lacking the N-terminal region, *in vitro* splicing reactions were assembled on Minx-P1R and supplemented with SDE2 lacking residues 78-95 at a final concentration of ∼ 1.2 uM. In this case no RNase H digestion was performed, and spliceosomes were captured via MS2 affinity, and then concentrated by sequential Strep-tactin affinity. Peak fractions were dialyzed as above and used directly for cryo-EM grid preparation. For some preps a quick concentration step after dialysis was performed by spinning at 6000 rpm for 3-6 minutes in an Amicon concentrator with 100 kDa MWCO. All samples were used immediately for EM sample preparation.

### Graphene grids preparation

The home-made graphene films were synthesized on 25 μm-thick copper foil (Alfa Aesar) by ambient-pressure chemical vapor deposition at 1070 °C, using methane as the carbon source, and subsequently transferred onto EM grids according to a previously reported method ^64^.

### Electron microscopy imaging

For cryo-EM analysis of the b-globin P complex, the sample was applied to R2/2 holey carbon grids (Quantifoil) coated with a ∼5-7 nm homemade carbon film. Grids were glow discharged for 15 s in an Edwards or Harrick plasma cleaner before application of 3 mL sample, then incubated for 25 s before blotting for 2.5-3.5 s (using in-house prepared blotting paper) and vitrification by plunging into liquid ethane using an FEI Vitrobot MKIV at 100% humidity and 8 °C. For Minx-P1R treated with the RH5 oligo the sample was applied to R1.2/1.3 holey carbon grids (Quantifoil) coated with homemade graphene film, while for the Minx-P1R treated with the RH3 oligo the sample was applied to commercial R2/2 holey carbon grids overlaid with a 2 nm carbon film (Quantifoil). For b-globin, grids were imaged at the MRC LMB and at Diamond eBIC in three separate sessions on FEI Titan Krios transmission electron microscopes operated in EFTEM mode at 300 kV using either a Gatan K2 Summit or a Gatan K3 direct electron detector and a GIF Quantum energy filter (slit width 20 eV); the camera was operated in counting mode with a total exposure time of 10 s fractionated into 40 frames, and a total dose of 45-60 e^-^ Å^-2^ per movie. For the Minx-P1R complexes, grids were imaged at Oxford COSMIC on an FEI G2 Titan Krios transmission electron microscope operated in EFTEM mode at 300 kV using a Gatan K3 direct electron detector and a GIF Quantum energy filter (slit width 20 eV); the camera was operated in counting mode with a total exposure time of 2 s fractionated into 40 frames, a dose rate of approximately 11.5 e^-^ pixel^-1^ s^-1^ and a total dose of 39-42 e^-^ Å^-2^ per movie.

### Image processing

All processing was performed using Relion 5.0^65,66^. Movies were corrected for movement using MotionCor2^67,68^, applying 5×5 patching without dose-weighting, and using the calibrated pixel sizes from specific microscopes outlined in figs. S1, S3 and S7. CTF parameters were estimated using Relion’s own implementation of CtfFind.

Specific step-by-step processing is outlined in figs. S1, S3 and S7. For all datasets particle picking was performed in Relion 5.0 with references obtained from our previous P complex structure^48^. For initial 2D and 3D classification particles were downscaled by a factor of 4 for the b-globin datasets and by a factor of 5 for the Minx-P1R dataset. Particles from the good 3D classes were then re-extracted and scaled to 1.2 Å/pixel (b-globin) or 1.66 Å/pixel (Minx-P1R) to allow subsequent merging of particles from multiple datasets. Following 3D refinement, Bayesian polishing and CTF refinement (per-particle defocus, anisotropic magnification, beam tilt, trefoil, and fourth-order aberrations) were performed to improve signal-to-noise ratio and accuracy of the reconstructions. Due to dataset heterogeneity, the b-globin DLS dataset was removed for the final classification and refinement steps and is not included in the final map.

For global 3D classification T=4 was generally used. To improve the map quality for the peripheral regions, particles from all datasets were merged, refined and then subjected to focused classification in Relion 5.0, without signal subtraction with soft masks around specific parts of the complex^69^ (figs. S1, S3 and S7). The reported resolutions were rounded to a single decimal and are based on the gold-standard Fourier Shell Correlation, as described ^70,71^. For the Minx-P1R datasets, note that since the images were binned to 1.66 Å/pixel during initial processing to allow for affordable computational costs on our cluster, the final reported resolutions from both Relion and Phenix are inherently limited to 3.32 Å by the Nyquist limit. See all FSC curves in figs. S1, S3 and S7.

### Model building and refinement

The models were built using Coot^72^ and ISOLDE^73^ starting with PDB 6QDV as an initial model for the core and the periphery^48^. The core was manually corrected in Coot and ISOLDE and then refined in PHENIX real_space_refine^74,75^ using one macro-cycle. Refinement was performed with hydrogens present, which were removed prior to deposition. Figures were generated using UCSF ChimeraX^76^ and PyMol.

### Declaration of generative AI and AI-assisted technologies in the manuscript preparation process

During the preparation of this work, the authors used Google Gemini and ChatGPT for language editing to improve clarity. The authors conceived all interpretations, reviewed all content, independently edited the final text, and take full responsibility for the content of the published article.

## Notes

### Competing Interest Statement

The authors have declared no competing interest.

