## Supplemental Materials for "Structural basis for alternative 3′ splice site selection in the human spliceosome active center"

#### Supplementary Notes

##### Cooperation of SDE2 and FAM32A during catalytic progression (Figures S3 and S4)

While SDE2, FAM32A, and Prp18 all participate in the exon-ligation phase, our structural and biochemical data suggest that they act in a defined sequential pathway to modulate the stability of the remodelled C* conformation and establish a docking-competent configuration of the active center (Fig. S4A).

Notably, SDE2 has both independent and cooperative roles during this process. Our data support a model wherein SDE2 likely acts first to stabilize the repositioned branch helix, which becomes flexible following Prp16-mediated remodeling[20, 23], and requires specific stabilization by human exon-ligation factors. Through this mechanism, SDE2 would increase the probability of the spliceosome reaching the productive, factor-engaged C*/P-like state. This model is supported by the observation that deletion of the SDE2 N-terminus (SDE2 ΔN) fundamentally changes the structural distribution of the spliceosome population: 66% of particles accumulate in a poorly engaged C*-like state lacking stable exon-ligation factors, while only 33% manage to reach the factor-engaged P-like state (Fig. 2C). By contrast, deletion of the FAM32A C-terminus did not cause a comparable shift in equilibrium, as other exon-ligation factors can still associate with the FAM32A ΔC complexes (Fig. S9D). These data are therefore consistent with a model in which SDE2 modulates the C-to-C* conformational equilibrium in a manner that is at least partly independent of the FAM32A C-terminus.

Within the productive, docking-competent C* state, FAM32A and Prp18 act in cooperation with SDE2 to stabilize the docked 3′-ss. Notably, unlike the SDE2 N-terminal deletion, truncation of the FAM32A C-terminus (FAM32A ΔC) does not generally cause a massive accumulation of lariat intermediates or strongly block exon ligation on optimal substrates (Fig. S4B, Fig. 3C). Instead, as visualized in our focused classification of the Minx-P1R structures (Fig. S9), the FAM32A C-terminus is specifically required to stabilize the docking of the 3′-ss and the surrounding mRNA junction within the active center, working alongside the SDE2 K78 residue to promote pairing of the 5′-ss and 3′-ss.

This specific requirement for FAM32A in stabilizing a closed C* state during 3′-ss docking is further illustrated by structural analysis of the active site in the two SDE2 ΔN cryo-EM classes. In class I, where FAM32A is absent, the pairing between the 3′-ss and 5′-ss is not yet ordered, and thus there is no clear density for a newly-formed mRNA bond (Fig. S3F). By contrast, in class II, where FAM32A is bound, the 3′-ss and 5′-ss are paired. Moreover, class II has clear density for both a newly-formed mRNA exon-exon bond, and for a docked and uncleaved 3′-ss / 3′-exon bond (Fig. S3F), consistent with a mixture of pre- and post-catalytic C*/P complexes.

Finally, the enhanced exon-ligation defect observed when combining SDE2 depletion with the FAM32A ΔC mutant (Fig. S4, B and C) implies that these factors affect distinct but connected steps during formation of a catalytically competent C* complex. We propose that this additive effect reflects a compounding kinetic barrier across two sequential probabilistic steps in a single pathway from the open to the closed C* complex (Fig. S4A): first, formation of a productive factor-engaged state, termed here C* "semi-open," which depends strongly on SDE2-dependent stabilization of the branch helix; and, second, stable 3′-ss docking to convert semi-open C* into closed C*, where FAM32A, Prp18, and SDE2 K78 act cooperatively. Thus, spliceosomes that manage to overcome the initial defect imposed by impaired SDE2 function (the 33% Class II population) remain vulnerable to failure to progress through the 3′-ss docking step when the FAM32A C-terminal anchor is absent. This model provides a mechanistic explanation for the enhanced lariat-intermediate accumulation observed in extract-based splicing assays.

##### Transcript-dependent engagement of Prp18 and 3′-ss docking (Figure S5)

In the human spliceosome, Prp18 engages the spliceosome core, an association not previously detected in any human complex assembled on viral Minx pre-mRNA. In the β-globin spliceosome, Prp18 binds the Prp8 RNase H domain, while its conserved loop abuts the 3′-ss and cooperates with the Prp8 α-finger to create a channel for 3′-ss docking. Pre-mRNA sequences may contribute to Prp18 engagement with the human C* and P spliceosomes. The MINX and β-globin substrates differ in their branch point to 3′-ss distance and their potential for 5′-exon to U5 loop I pairing (Fig. S5A). Strong 5′-exon pairing to U5 snRNA loop I in Minx correlates with low, salt-sensitive association of Prp18, whereas weaker pairing, as observed for β-globin, increases stable, salt-resistant Prp18 association (Fig. S5E). Cryo-EM reconstructions confirm that Prp18 binds stably only in β-globin P complexes, where weakened 5′-exon positioning correlates with complete ordering of the FAM32A C-terminus. Prp18 actively stabilizes the correct juxtaposition of the 5′-exon nucleophile with the docked 3′-ss, a role that may be particularly important for pre-mRNAs with weaker splice-site interactions with the spliceosome core.

### Supplementary Figures and Tables

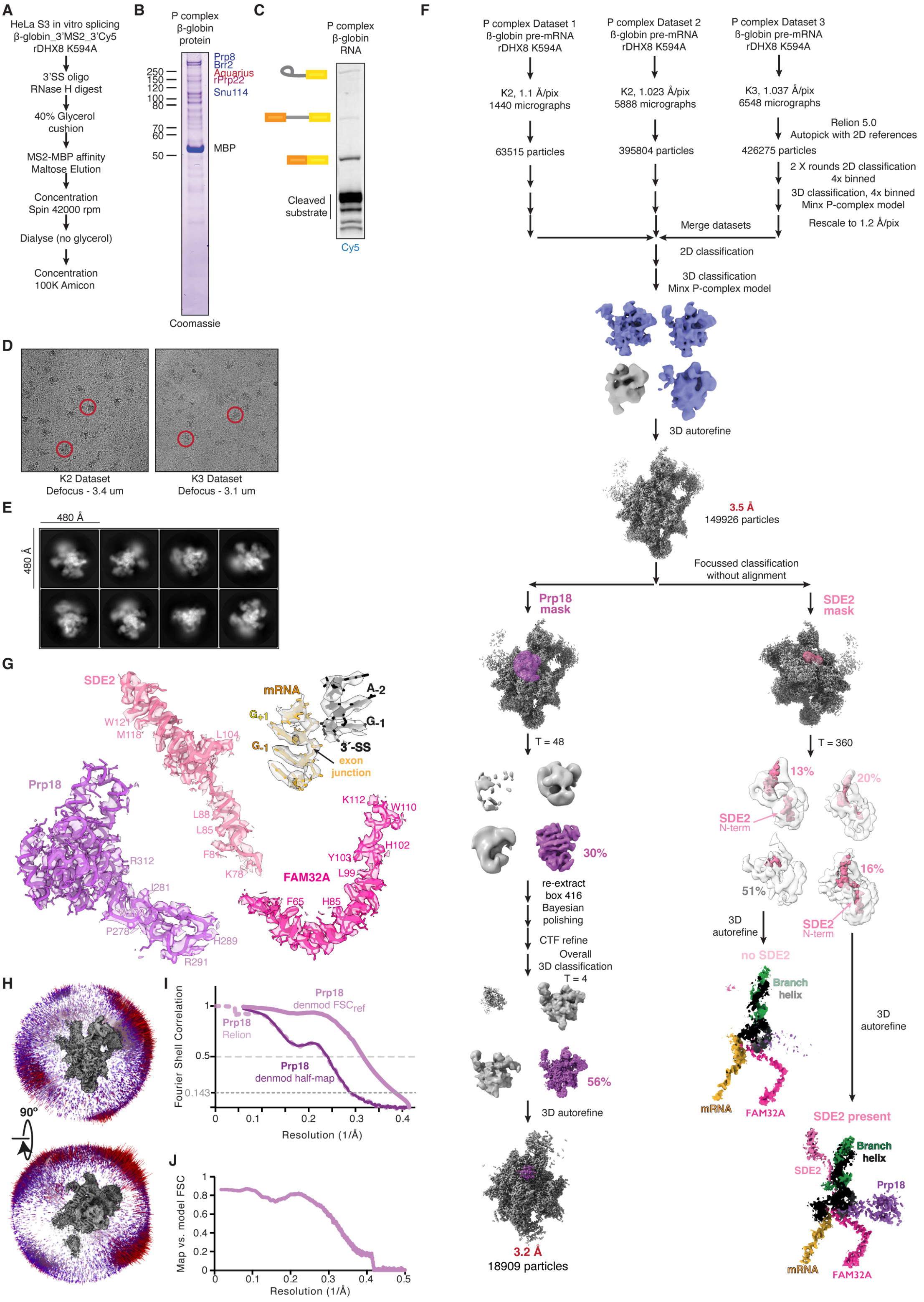


**Figure S1. Cryo-EM processing and structure determination of the human β-globin P-complex spliceosome.** **(A)** Schematic of MS2 affinity purification of spliceosomes assembled on the β-globin pre-mRNA substrate. **(B and C)** Coomassie-stained SDS–PAGE **(B)** and RNA gel **(C)** showing protein and RNA composition of purified spliceosomes. **(D)** Representative cryo-EM micrographs showing well-dispersed particles (red circles). **(E)** 2D class averages reveal characteristic P-complex projections. **(F)** Relion data-processing workflow. Focused classification with masks on Prp18 and SDE2 revealed distinct states and improved resolution of specific factors. **(G)** Cryo-EM density and model for SDE2, Prp18, FAM32A, the mRNA, and the 3′-ss. **(H)** Representative angular distribution for the cryo-EM map of the β-globin P-complex spliceosome (refined Prp18-focused map). **(I and J)** Fourier shell correlation (FSC) for the refined maps **(I)** and the overall model versus the composite map **(J)**; denmod, density-modified maps.

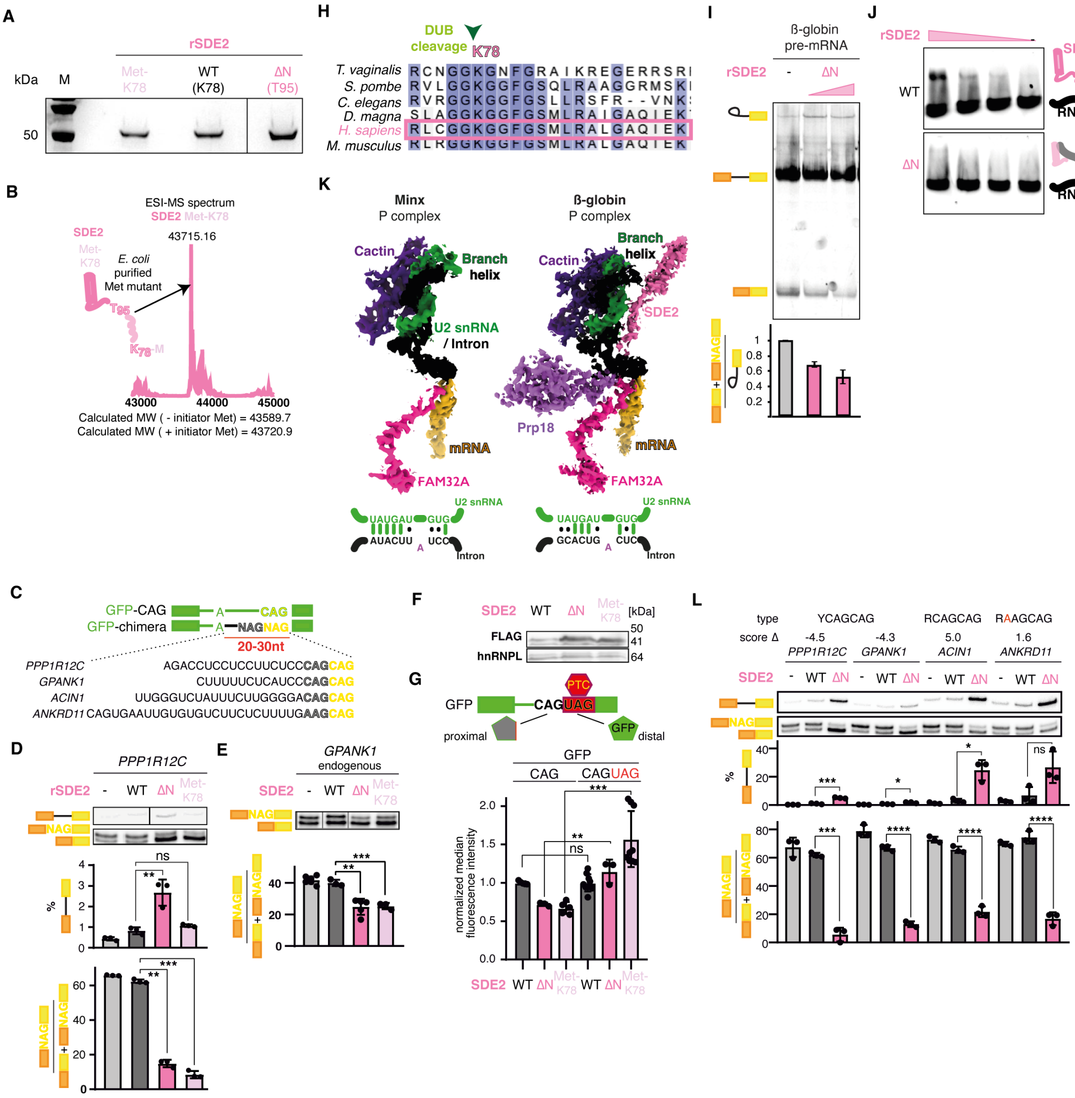

**Figure S2. Functional analysis of SDE2 during exon ligation and 3′-ss selection.** **(A and B)** Coomassie-stained SDS-PAGE **(A)** and mass spectrometry validation **(B)** of recombinant SDE2 proteins: wild-type (WT K78), N-terminal deletion (ΔN T95), and methionine-insertion mutant (Met-K78). **(C)** Structure of heterologous minigenes used for *in vivo* assays, capturing diverse PY tracts and NAGNAG sequences. **(D and E)** SDE2 mutations shift alternative 3′-ss selection *in vivo*. Representative RT-PCR and quantification for the *PPP1R12C* minigene **(D)** and the endogenous *GPANK1* intron **(E)**. The SDE2 Met-K78 mutant acts purely as a regulatory switch for distal 3′-ss usage without causing intron retention (IR), whereas SDE2 ΔN both shifts splicing and strongly induces IR (quantified in the upper graph of **D**). **(F and G)** SDE2 disruption shifts NAGNAG selection in a GFP reporter *in vivo*, activating distal GFP expression via skipping of the UAG stop codon. Protein expression of the different SDE2 variants was verified by Western blot **(F)** alongside FACS quantification of normalized GFP fluorescence **(G)**. **(H)** Conservation of the SDE2 N-terminus across eukaryotes. **(I)** *In vitro* splicing of β-globin pre-mRNA confirms SDE2 ΔN impairs overall exon-ligation efficiency. **(J)** EMSA demonstrates the SDE2 N-terminus promotes RNA binding *in vitro*. **(K)** Comparison of cryo-EM maps for P-complex spliceosomes assembled on viral (MINX, EMDB-4525) versus β-globin (current study) substrates reveals distinct engagement of the SDE2 N-terminus and branch helix. **(L)** RT-PCR of diverse NAGNAG minigenes confirms SDE2 ΔN broadly shifts splicing to distal acceptors and induces IR across multiple sequence contexts. For panels **D**, **E**, **G**, and **H**, data are represented as mean ± SD of 3–6 independent experiments. Statistical significance calculated by Student's t-test: *, $p < 0.05$; **, $p < 0.01$; ***, $p < 0.001$; ****, $p < 0.0001$; ns, not significant.

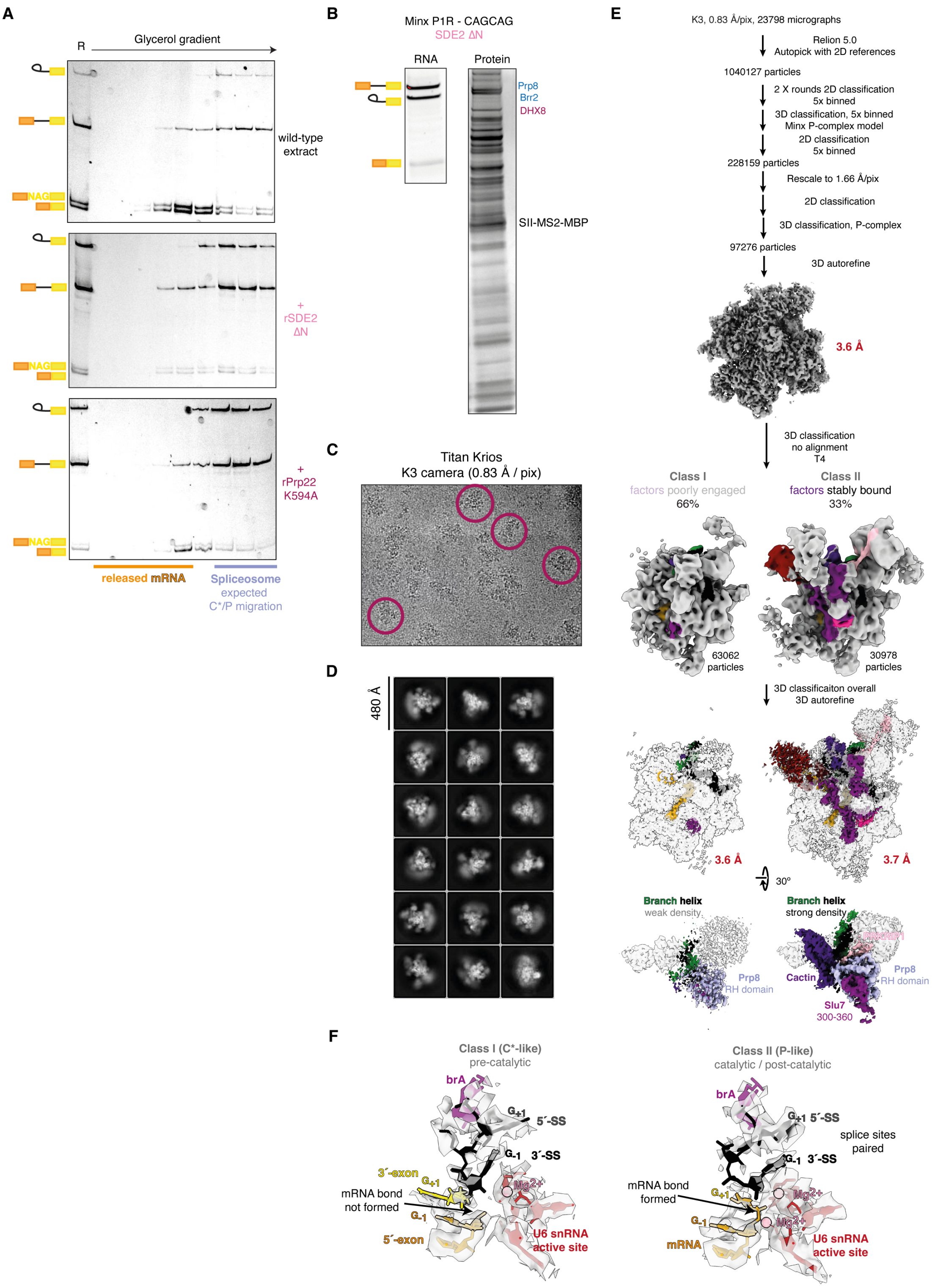


**Figure S3. Cryo-EM processing of SDE2 ΔN spliceosomes and functional analysis of SDE2-mediated C* complex stabilization.** **(A)** Glycerol-gradient sedimentation of *in vitro* splicing reactions reveals that SDE2 ΔN leads to accumulation of spliceosomes in fractions corresponding to stalled pre-catalytic C*-like particles, consistent with impaired catalytic turnover. For comparison, accumulation of C*/P complexes is shown upon blocking mRNA release with Prp22 K594A. **(B)** RNA and protein composition of purified C*/P spliceosomes assembled on the Minx-P1R substrate in the presence of SDE2 ΔN. **(C and D)** Representative cryo-EM micrograph **(C)** and 2D class averages **(D)** of the SDE2 ΔN sample. **(E)** Data-processing workflow for the SDE2 ΔN dataset. 3D classification separates particles into Class I (66%), lacking stable engagement of exon-ligation factors and exhibiting weak branch helix density, and Class II (33%), showing stably bound factors and strong branch helix density. **(F)** Cryo-EM density and models for the core of the class I C*-like, pre-catalytic complexes, and of the class II P-like catalytic/ post-catalytic spliceosomes observed in the presence of SDE2 ΔN.

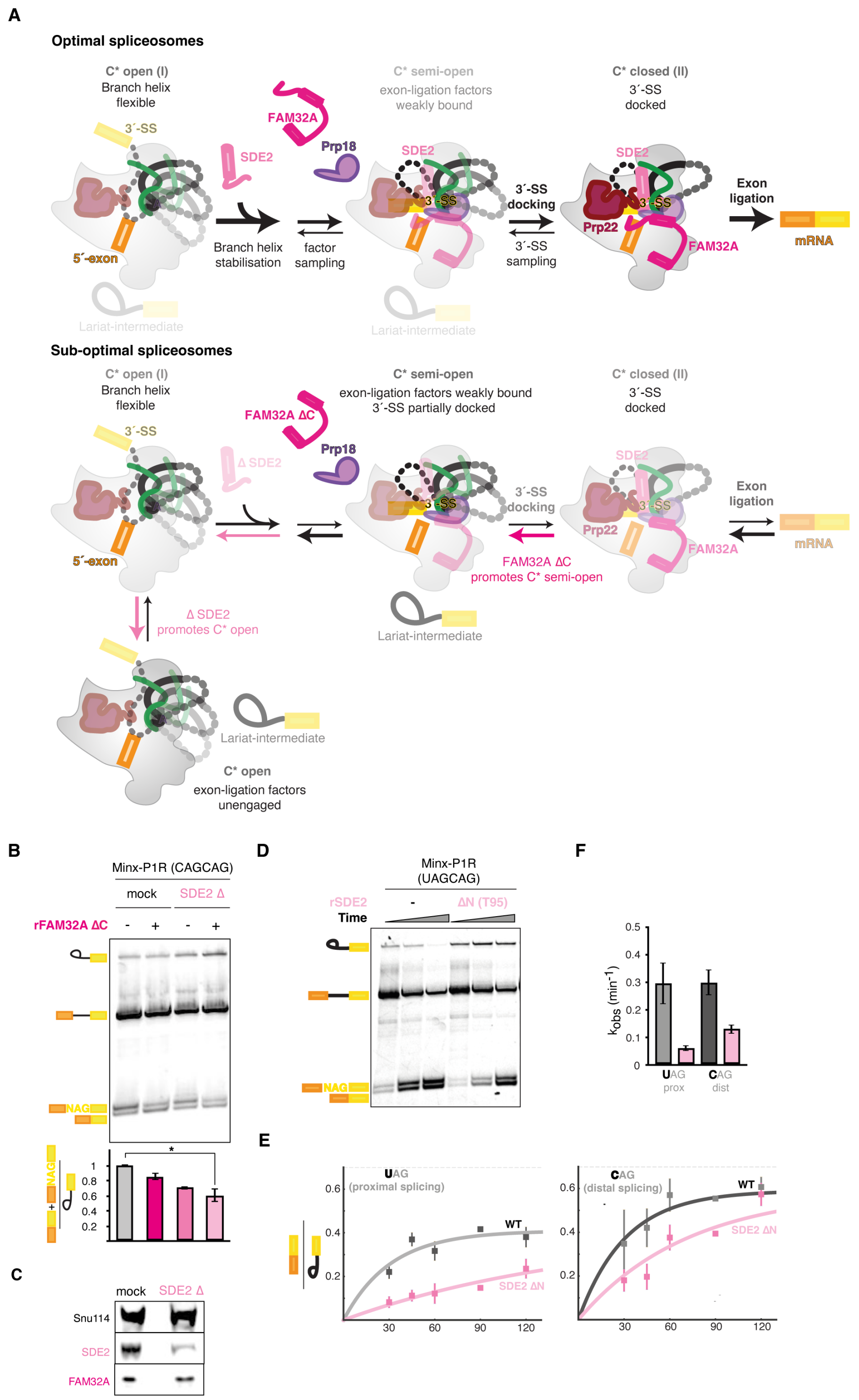


**Figure S4. SDE2 and FAM32A modulate conformational equilibrium of the spliceosome during 3′-ss docking and catalytic progression. (A)** Schematic depicting the structural equilibrium of the spliceosome active center during the transition to exon ligation. In optimal spliceosomes (top), SDE2 stabilizes the branch helix to shift the landscape from a flexible C* open state toward a factor-sampling C* semi-open state. Subsequently, FAM32A and Prp18 cooperate with SDE2 to stabilize 3′-ss docking and form the catalytically competent C* closed state. In sub-optimal spliceosomes (bottom), loss of SDE2 (Δ SDE2) shifts the structural equilibrium backwards to the unengaged C* open state, resulting in lariat-intermediate accumulation. Conversely, truncation of the FAM32A C-terminus (FAM32A ΔC) specifically destabilizes the 3′-ss, shifting the equilibrium back toward the partially docked C* semi-open state. **(B and C)** Depletion of SDE2 shifts 3′-ss use in a modified MINX substrate with the *PPP1R12C* CAGCAG site (Minx-P1R), and the addition of recombinant FAM32A ΔC exacerbates this effect. *In vitro* splicing and quantification **(B)** and Western blot verifying specific depletion of SDE2 **(C)** are shown. Data are represented as mean ± SD of 3 independent experiments, from two independent depletions; *, $p<0.05$, paired t-test. **(D to F)** Deletion of the SDE2 N-terminus slows the kinetics of exon ligation. *In vitro* splicing time course **(D)** and kinetic quantification **(E)** for proximal (UAG) and distal (CAG) splice sites on a Minx-P1R substrate. Calculated $k_{obs}$ values **(F)** demonstrate that SDE2 ΔN significantly impairs the rate of exon ligation at both sites.

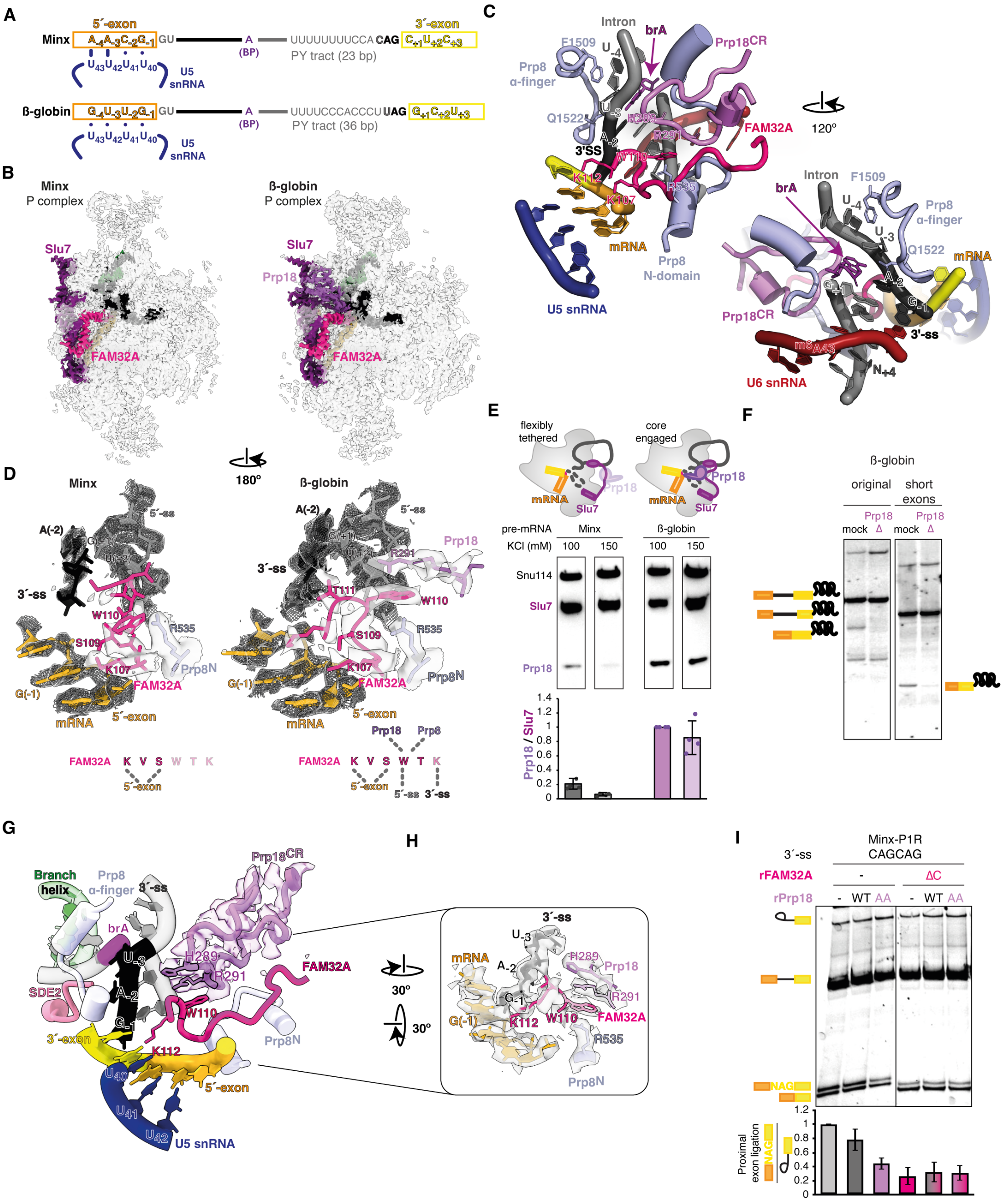

**Figure S5. Prp18 engages the spliceosome core in a transcript-dependent manner to promote 3′-ss docking.** **(A)** MINX and β-globin substrates differ in BP to 3′-ss distance and 5′-exon/U5 loop I pairing potential. **(B to D)** Cryo-EM reconstructions show that Prp18 binds stably only in β-globin P complexes, where weakened 5′-exon positioning correlates with complete ordering of the FAM32A C-terminus **(B)**. Structural details **(C)** reveal that Prp18, the Prp8 α-finger, the FAM32A WTK motif, and the Prp8 N-domain form a channel that guides docking and stabilizes the 3′-ss. In the Minx structure **(D)**, there is no strongly ordered density for FAM32A W110-K112 (the model is obtained by restrained refinement but is out of density); by contrast, in β-globin, ordered density is clearly observed for W110-K112. **(E)** Western blots confirm transcript-specific recruitment of Prp18, with higher Prp18:Slu7 ratios on β-globin spliceosomes. Prp18 levels were normalized to Slu7 for individual conditions, and relative Prp18 levels were then further normalized to the Prp18 signal for β-globin spliceosomes washed with 100 mM KCl. Data are shown as mean ± SD of 3–4 independent purifications. **(F)** Prp18 depletion compromises exon ligation of β-globin constructs. Structural studies were performed with a modified β-globin pre-mRNA where the flanking exons were truncated to 45 nucleotides (short exons) to match the length of the Minx exons. For comparison, depletion of Prp18 also impairs splicing of a β-globin pre-mRNA containing the original full-length flanking exons. **(G to I)** Prp18 cooperates with FAM32A to modulate alternative 3′-ss selection. Close-up of the docked 3′-ss shows the Prp18 conserved loop (CR) bridges the docked 3′-ss to FAM32A **(G)**, whose WTK motif is stabilized by Prp18 R291 and juxtaposes the 5′-exon and 3′-ss **(H)**. **(I)** Prp18 and FAM32A modulate NAGNAG 3′-ss use *in vitro*. Bar graph quantifies proximal exon-ligation efficiency, normalized to no protein added. Data are represented as mean ± SD (n = 4–6).

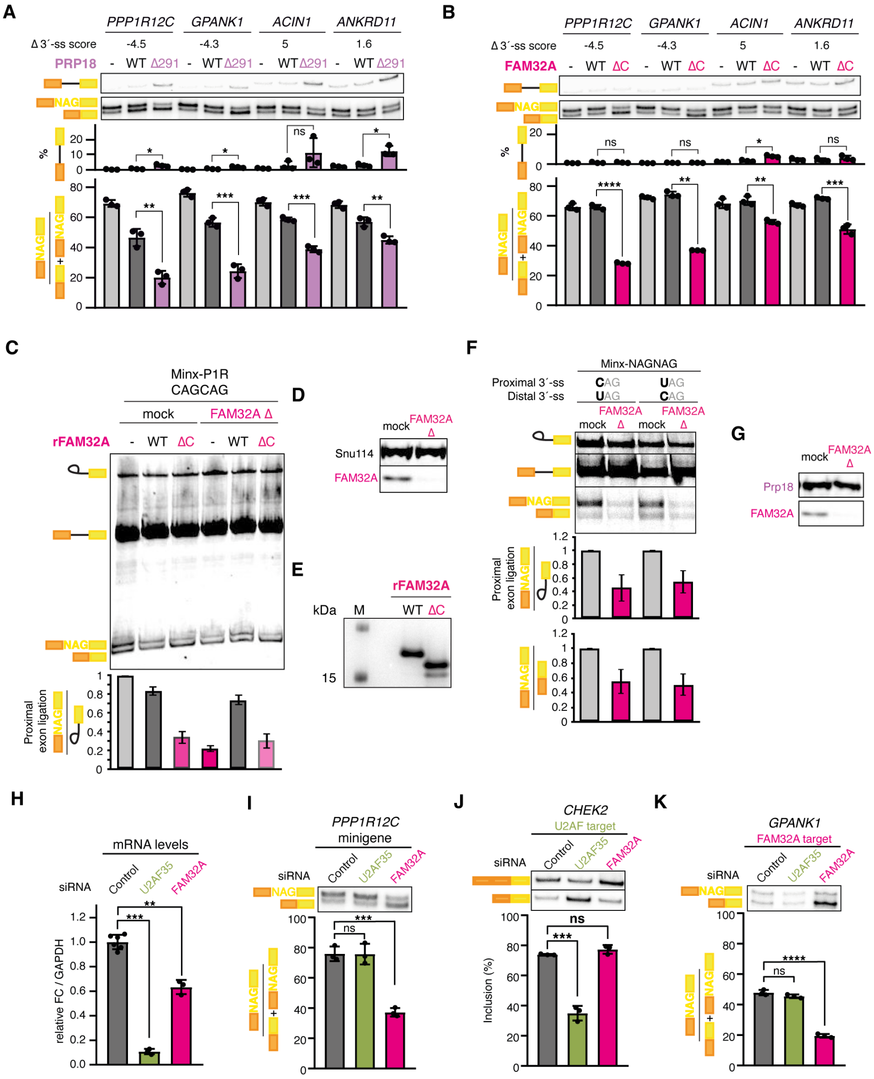


**Figure S6. Prp18 and FAM32A cooperate to promote proximal NAGNAG 3′-ss use during catalysis.** (A and **B)** Prp18 Δ291 **(A)** and FAM32A ΔC **(B)** decrease proximal NAGNAG 3′-ss usage in selected minigenes, depending on the relative strength of competing 3′-ss. Negative scores reflect a stronger distal site score and stronger regulation, calculated as proximal score minus distal score. **(C)** FAM32A acts specifically during the catalysis of exon ligation. *In vitro* splicing assays in control or FAM32A-depleted extracts complemented with recombinant FAM32A WT or ΔC. **(D)** Western blot analysis confirming selective FAM32A depletion. **(E)** SDS-PAGE of purified recombinant FAM32A proteins used in the biochemical assays. **(F to G)** *In vitro* splicing of Minx-NAGNAG substrates with different combinations of proximal and distal CAG/UAG 3′-ss. Notably, introducing a NAGNAG site into the classic Minx substrate, which possesses a standard polypyrimidine tract, demonstrates that the tandem 3′-ss itself is sufficient to induce FAM32A-dependent regulation. Depletion of FAM32A **(F)** strongly impairs proximal 3′-ss usage. Bar graphs quantify proximal exon-ligation efficiency (top) and relative use of proximal versus distal sites (bottom), normalized to mock depletion. Western blots validate selective depletion of FAM32A **(G)** from splicing extracts. **(H** to **K)** FAM32A acts independently of U2AF to modulate 3′-ss selection. Knockdown efficiency of U2AF35 and FAM32A was quantified by RT-qPCR relative to GAPDH **(H)**. The chimeric PPP1R12C minigene **(I)** and the endogenous GPANK1 **(J)** FAM32A targets are unaffected by U2AF depletion, which only causes exon skipping for the endogenous CHEK2 U2AF target **(K)**. For all panels, data are represented as mean ± SD of 3–4 independent experiments. Statistical significance was calculated by Student's t-test: *, $p < 0.05$; **, $p < 0.01$; ***, $p < 0.001$; ****, $p < 0.0001$; ns, not significant.

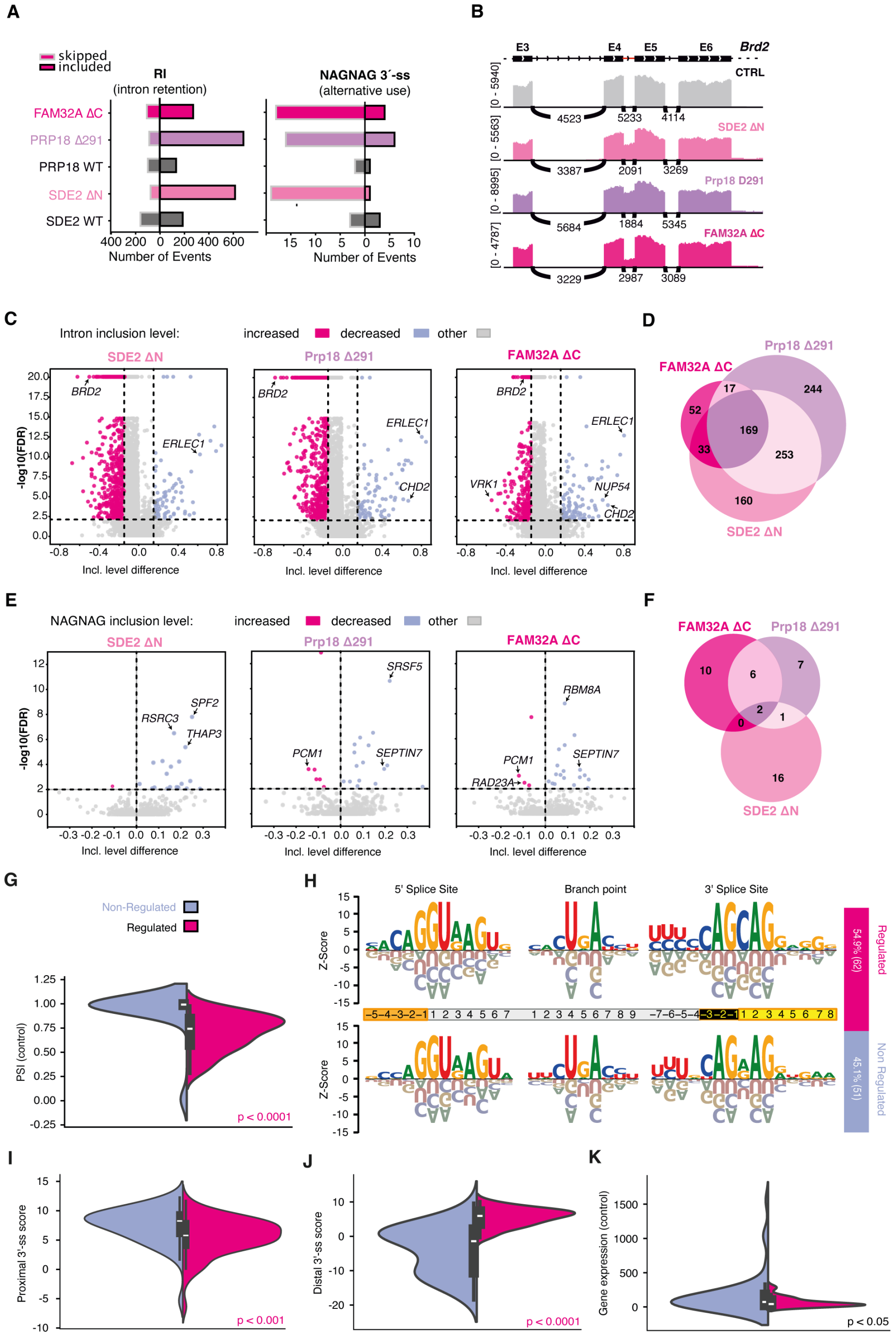


**Figure S7. Genome-wide effects of C* exon-ligation factor mutants and cis-features of FAM32A-regulated introns.** **(A)** Prp18 and SDE2 mutants cause genome-wide intron retention (RI), but FAM32A has a more specific role in alternative NAGNAG 3′-ss selection. The number of significantly altered events detected by RNA-seq is shown. **(B)** Prp18, SDE2 and FAM32A have distinct effects on intron retention *in vivo*. Genome-browser views of the *BRD2* locus show accumulation of intronic reads for intron 4 (but not introns 3 and 5) in response to overexpression of specific factor mutants. CTRL, control non-transfected cells. **(C)** Volcano plots for intron retention events (inclusion-level difference versus –log10 FDR) upon overexpression of SDE2 ΔN, Prp18 Δ291, and FAM32A ΔC relative to CTRL; significantly increased and decreased events, along with select transcripts, are highlighted. **(D)** Venn diagram showing overlap of introns with significantly increased retention. **(E)** Volcano plots of NAGNAG inclusion-level changes in the same datasets. **(F)** Venn diagram showing overlap of NAGNAG exons with increased distal-site usage among the three mutants. (**G to K**) FAM32A regulates a specific subset of NAGNAG sites. Target introns are based on previous work using siRNA against FAM32A. **(G)** Violin plots of basal PSI (control) for FAM32A-regulated versus non-regulated NAGNAG 3′-ss. **(H)** Z-score sequence logos for the 5′-ss, branch point, and 3′-ss regions of FAM32A-regulated (top) and non-regulated (bottom) 3′-ss. Positions used for machine learning are highlighted in the middle. Note that FAM32A regulates a specific set of target introns, enriched for a strong distal CAG motif and a weak proximal -4 position (C/U). **(I and J)** Distributions of proximal **(I)** and distal **(J)** 3′-ss strength scores for FAM32A-regulated versus non-regulated introns. **(K)** Transcript levels for endogenous genes for regulated and non-regulated introns. For **G**, **I**, **J**, and **K**, statistical significance was determined by Mann–Whitney U test.

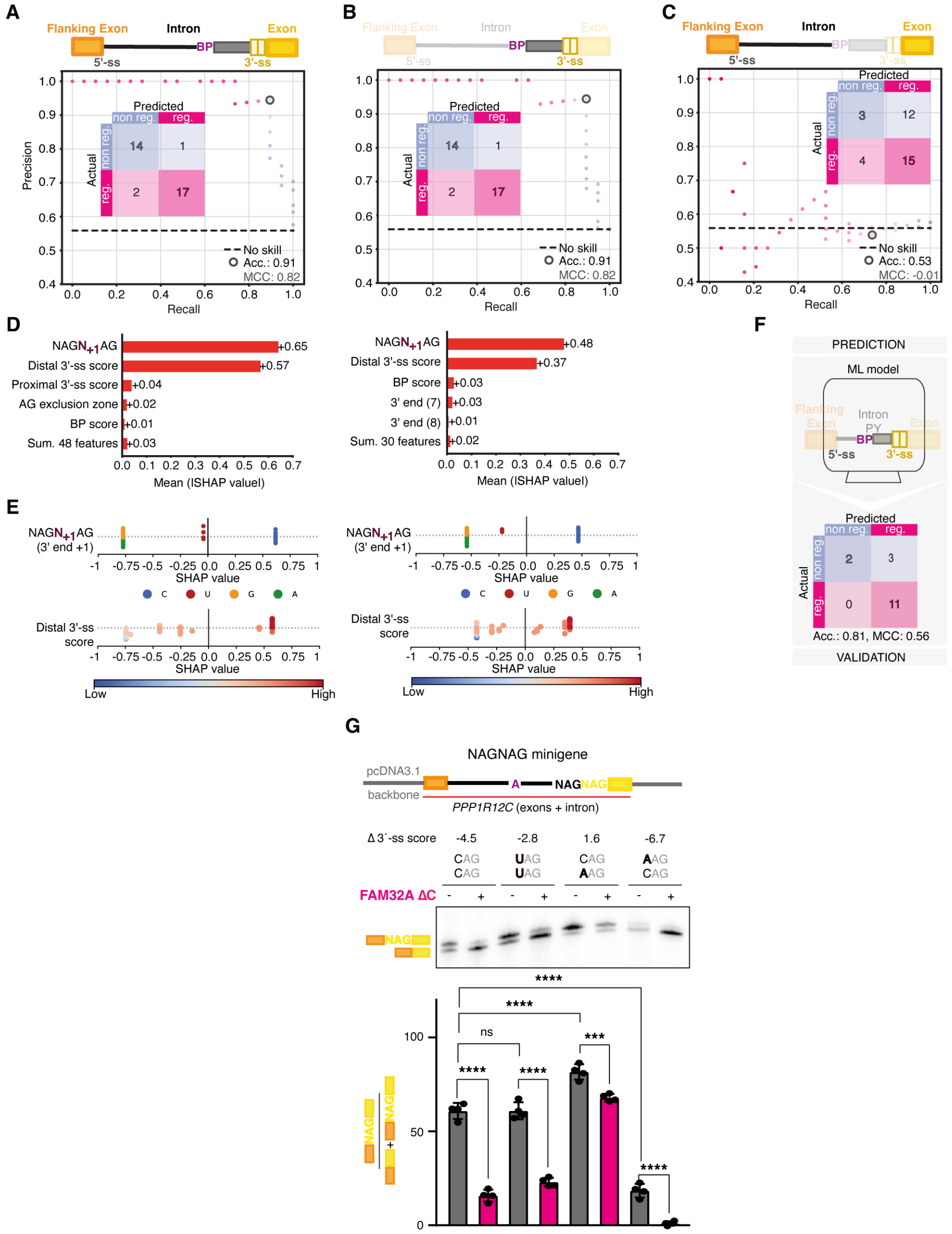


**Figure S8. Machine learning prediction and experimental validation of TASS regulation by exon-ligation factors.** **(A to C)** Precision–recall performance of classifiers trained on distinct feature sets. Schematics above the plots indicate included regions (shading). Shown are a full *cis*-feature model **(A)**, a reduced feature model retaining key BP to 3′-ss features **(B)**, and a control model lacking only the 3′-ss context **(C)**. Insets show confusion matrices for the operating threshold highlighted on the curves; Matthew's correlation coefficient (MCC) is indicated. While models containing all features or only 3′-ss features perform comparably, a model without 3′-ss features shows no predictive skill. **(D)** Global feature importance ranked by mean absolute SHAP value for the full model (top, from **A**) and reduced model (bottom, from **B**), highlighting dominant contributions from 3′-end composition (position number relative to start of the distal NAG; N+1 = -3 nucleotide of the distal 3′-ss). Predictive power of both models is based on the +1 position identity and the distal 3′-ss score. **(E)** SHAP summary plots showing the direction and magnitude of feature effects across events for the full model (left) and reduced model (right); points are colored by nucleotide identity at the 3′-end position or relative distal 3′-ss score strength. FAM32A target introns are strongly enriched for C at the +1 position and a high distal 3′-ss score. The reduced model was used for all downstream analyses to validate predictions using minigenes containing the BP-to-3′-ss sequences of interest. **(F)** Evaluation of the reduced model predictions using the responsiveness of minigene mutations to FAM32A ΔC (from **G** and Figure 3D), shown as a confusion matrix with overall accuracy and MCC. **(G)** FAM32A ΔC overexpression decreases proximal NAGNAG 3′-ss usage *in vivo* across selected minigenes. Variations at the −3 position result in different strengths of the competing splice sites and influence the effect of FAM32A ΔC. A minigene system containing the entire *PPP1R12C* NAGNAG intron with surrounding exon sequences is shown. The calculated "Δ 3′-ss score" indicates the difference between proximal and distal 3′-ss scores. All variants contain a C at -4, and a weak proximal AAG requires FAM32A for proximal site usage. Data are represented as mean ± SD of 3 independent experiments. Statistical significance calculated by Student's t-test: ***, $p < 0.001$; ****, $p < 0.0001$; ns, not significant.

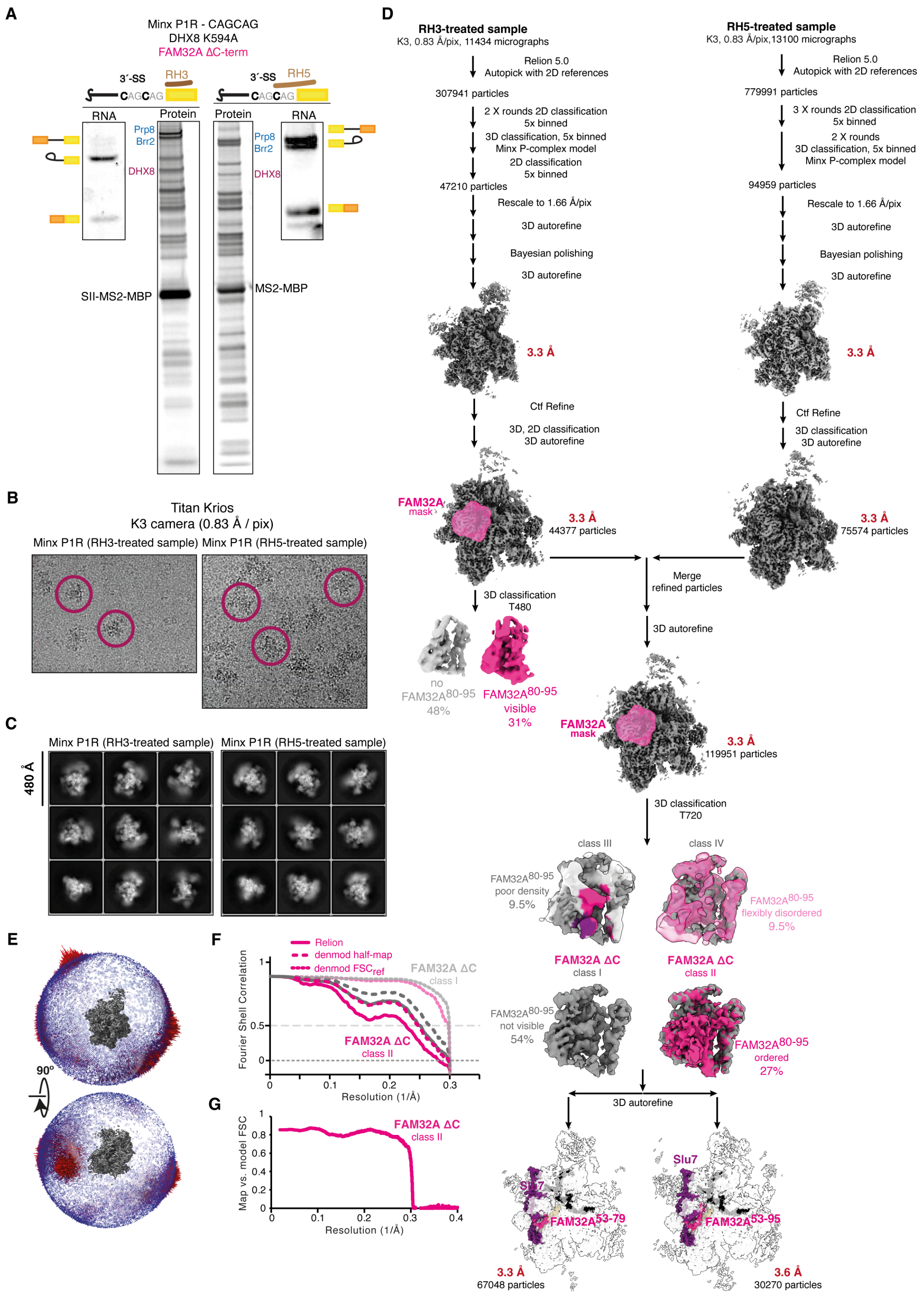


**Figure S9. Cryo-EM data collection and processing for Minx-P1R P-complex spliceosomes.** **(A)** SDS-PAGE of affinity-purified spliceosome complexes assembled on Minx-P1R pre-mRNA in the presence of FAM32A ΔC and treated with either the RH3 or RH5 oligo, showing associated RNA and protein components. Note that this procedure purifies only complexes with a partially protected/docked 3′-ss. **(B)** Representative cryo-EM micrographs recorded on a Titan Krios microscope for RH3- and RH5-treated samples. **(C)** Selected two-dimensional class averages of particle images from each dataset. **(D)** Single-particle analysis workflow for RH3- and RH5-treated data, including iterative 2D and 3D classification, Bayesian polishing, and focused classification with a FAM32A mask, yielding P-complex reconstructions in which the FAM32A 80–95 segment is absent or partially visible/ordered, together with particle numbers and final resolutions. **(E)** Representative angular distribution for the cryo-EM map of the FAM32A ΔC class II P-complex spliceosome. **(F and G)** Fourier shell correlation for the refined maps **(F)** and the model versus the class II map **(G).**

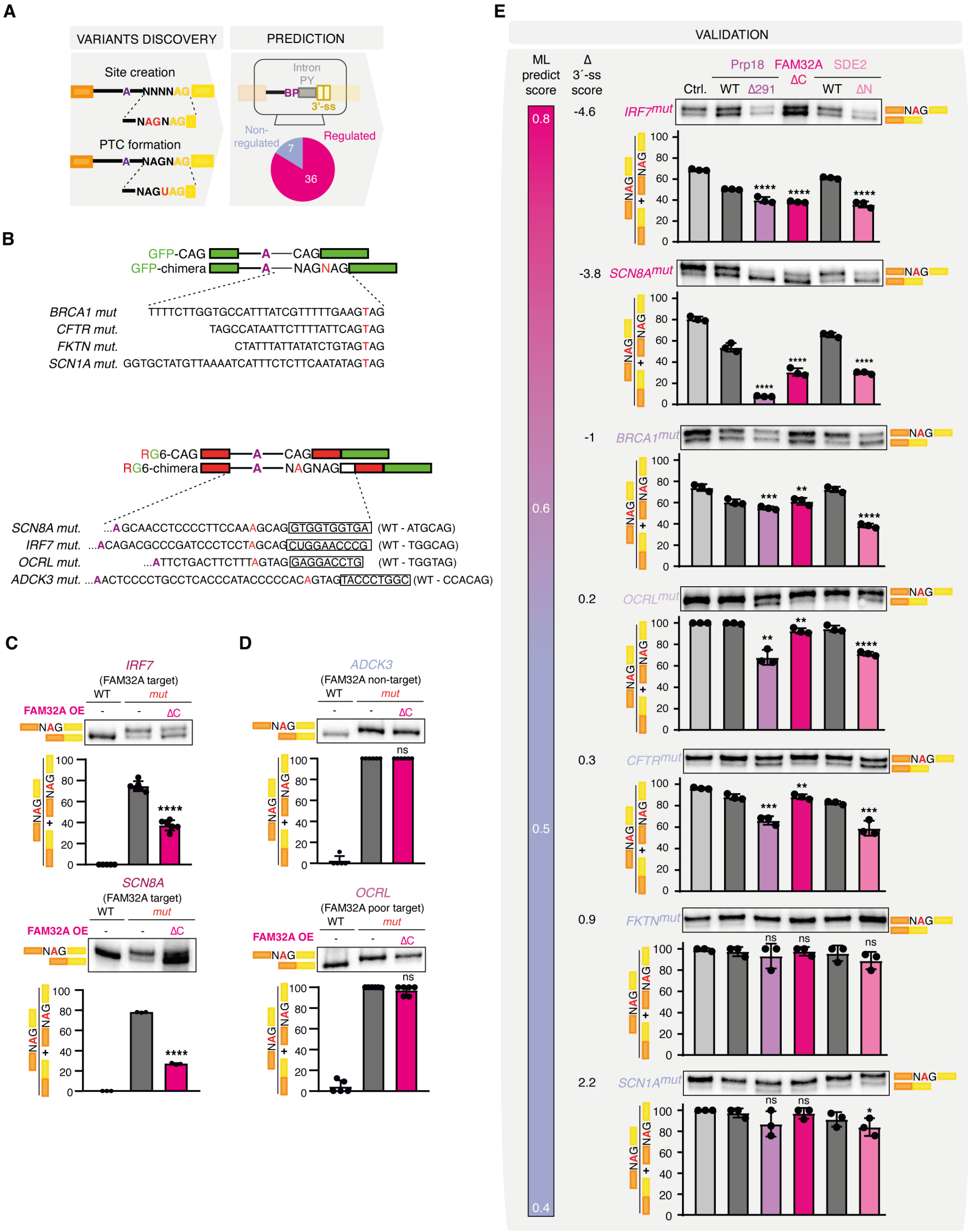


**Figure S10. Perturbation of exon-ligation factors restores accurate splicing for disease-relevant NAGNAG alleles.** **(A)** Regulation of disease-related targets by exon-ligation factors can be accurately predicted by machine learning. Selected single-nucleotide variants (SNVs) comprising AG-gain and PTC-creating mutations were predicted with our ML model; prediction rates are shown. **(B)** Sequence cartoon of minigenes used for *in vivo* splicing, harboring mutated splice sites of different identified disease targets. Upper panel: GFP-chimeras with inserted splice sites of PTC-gain targets. Lower panel: RG6-chimeras with varying complete splice sites including the first 10 nucleotides of the downstream exon of different AG-gain targets. The respective WT sequence is indicated on the right. **(C and D)** The emergence of a new proximal splice site leads to dominant usage of this site compared to the wild-type splice site. Modulation by FAM32A ΔC can restore wild-type distal splice site usage in predicted targets **(C)**, but not in predicted poor or non-targets **(D)**. The AG-gain variants in *IRF7* and *SCN8A* are in disease-relevant genes, but their direct disease relevance has not been confirmed. **(E)** Regulation of predicted disease-relevant targets correlates with the score difference between competing 3′-ss. The ML prediction probability is shown as a shaded bar graph (left), and the individual Δ 3′-ss scores are indicated; note that for both *BRCA1* and *CFTR*, the prediction probability is 0.52, but their ordering reflects the order of Δ 3′-ss scores. The actual probabilities for the other events are: *IRF7* 0.74, *SCN8A* 0.74, *OCRL* 0.59, *FKTN* 0.44, *SCN1A* 0.43. Representative RT-PCR gels and quantification of proximal isoform usage using minigenes with corresponding disease splice sites are shown (right); select targets from main Fig. 4 are reproduced for completeness. Skipping of a cryptic proximal splice site can be achieved by disrupting Prp18, FAM32A, or SDE2 function in specific targets, suggesting differential sensitivity to specific exon-ligation factors. For all panels, data are mean ± SD; points indicate biological replicates. Statistical significance is shown as indicated: *, $p < 0.05$; **, $p < 0.01$; ***, $p < 0.001$; ****, $p < 0.0001$; ns, not significant.

#### Table S1. Composition of purified P complexes[a]

| Sub-complexes | Protein | M.W. (kDa) | β-globin peptides | MINX P1R peptides | Modelled chain β-globin[b] | S. cerevisiae/ S. pombe names |
|---|---|---|---|---|---|---|
| U5 snRNP | PRPF8 | 274 | 70 | 63 | /A:25-2019 | Prp8/Spp42 |
| | Brr2 | 244.5 | 66 | 64 | Docked | Brr2/Brr2 |
| | 116K | 109.4 | 29 | 30 | /C:56-954 | Snu114/Cwf10 |
| | U540K / SNR40 | 39.3 | 6 | 9 | /N:49-357 | -/Cwf17 |
| | U5 snRNA | 39 | | | /5:9-70 | |
| Sm Ring | SmB | 24.6 | 3 | 5 | Docked | SmB/SmB |
| | SmD3 | 13.9 | 2 | 4 | Docked | SmD3/SmD3 |
| | SmD1 | 13.3 | - | 3 | Docked | SmD1/SmD1 |
| | SmD2 | 13.5 | 5 | 5 | Docked | SmD2/SmD2 |
| | SmF | 9.7 | 3 | 1 | Docked | SmF/SmF |
| | SmE | 10.8 | 2 | 1 | Docked | SmE/SmE |
| | SmG | 8.5 | 3 | 2 | Docked | SmG/SmG |
| U2 snRNP | U2-A' | 28.4 | 14 | 7 | Docked | Lea1/Lea1 |
| | U2-B'' | 25.5 | 5 | 4 | Docked | Msl1/Msl1 |
| | U2 snRNA | 63.9 | | | /2:1-53 | U2 snRNA |
| U6 | U6 snRNA | 36.1 | | | /6:1-5,15-97 | |
| NTC | PRPF19 | 55.2 | 6 | 9 | Docked | Prp19/Cwf8 |
| | SPF27/BCAS2 | 26.1 | 8 | 7 | Docked | Snt309/Cwf7 |
| | SYF1/XAB2 | 100 | 35 | 14 | Docked | Syf1/Cwf3 |
| | SYF2 | 28.7 | 6 | 7 | /y:118-243 | Syf2/Syf2 |
| | CRNKL1 | 100.5 | 38 | 16 | /S:126-437 | Clf1/Cwf4 |
| | CDC5L | 92.3 | 27 | 21 | /O:2-109,130-283 | Cef1/Cdc5 |
| NTR | SNW1/SKIP | 61.5 | 22 | 10 | /K:41-163,174-308 | Prp45/Prp45 |
| | PLRG1 | 57.2 | 8 | 8 | /J:185-504 | Prp46/ Prp5 |
| | RBM22 | 46.9 | 8 | 10 | /M:18-222 | Cwc2+Ecm2/ Cwf2+Cwf5 |
| | CWC15 | 26.6 | 4 | 1 | /P:2-75,187-197,209-229 | Cwc15/Cwf15 |
| | BUD31 | 17 | - | 5 | /L:4-144 | Bud31/Cwf14 |
| Splicing factors | CDC40 | 65.5 | 13 | 18 | /o:67-144,151-223,234-579 | Prp17/Prp17 |
| | SRRM2/SRm300 | 299.6 | 14 | 19 | /R:1-26 | Cwc21/Cwf21 |
| | CWC22 | 105.5 | 27 | 12 | /H:448-648 | Cwc22/Cwf22 |
| IBC protein | Aquarius | 171.3 | 29 | 36 | Docked | -/Cwf10 |
| | Isy1 | 33 | 9 | 3 | Not modelled | Isy1/Cwf12 |
| ATPases | DHX8 / hPrp22 | 139.3 | 35 | 36 | /V:1187-1218 | Prp22/Prp22 |
| | DDX41/Abstrakt | 70 | 34 | 27 | Not modelled | -/Dbj1 |
| Exon junction complex | eIF4AIII | 46.9 | 12 | 10 | Docked | Fal1/Fal1 |
| | RBM8A | 19.9 | - | 1 | Docked | -/Rbm8 |
| | MAGOH | 17.3 | 2 | 2 | Docked | -/Mnh1 |
| Exon-ligation factors | SLU7 | 68.4 | 19 | 11 | /c:25-200,262-363 | Slu7/Slu7 |
| | PRPF18 | 39.9 | 11 | 4 | /a:165-330 | PRP18/prp18 |
| | FAM32A | 13.2 | 3 | 2 | /G:53-112 | -/SPAC31G5.21 |
| | Cactin | 88.7 | 18 | 18 | /F:637-758 | -/Cay1 |
| | PRKRIP1 | 21 | 3 | 3 | /D:20-142 | -/SPBC16G5.11 |
| | PPIL1 | 18.2 | 5 | 2 | /i:4-166 | Cpr2//Cyp1 |
| | SDE2 | 49.7 | 11 | 2 | /z:78-126 | -/Sde2 |
| | NKAP | 46.9 | 4 | 3 | /Z:329-358 | -/Nkap1 |
| Other unmodelled proteins | PPIL3 | 18 | 5 | 7 | Not modelled | -/Cyp7 |
| | CCDC12 | 19 | 6 | 4 | Not modelled | -/Cdc12 |
| | CXorf56 | 26 | 7 | - | Not modelled | -/SPAC26F1.02c |
| | LENG1 | 31 | 4 | 2 | Not modelled | -/SPAC23H3.14c |
| | PPIE | 33 | 11 | 11 | Not modelled | -/Cyp9 |
| | NOSIP | 33 | 4 | 8 | Not modelled | -/SPBC3B9.01 |
| | C9orf78 | 34 | 4 | 6 | Not modelled | -/Tls1 |
| | YBX1 | 36 | 8 | 1 | Not modelled | -SPAC15A10.12 |
| | FAM50A | 40 | 7 | 6 | Not modelled | -/Xap5 |
| | FAM50B | 39 | 10 | 6 | Not modelled | -/- |
| | CWC27 | 54 | - | 2 | Not modelled | Cwc27/Cwc27 |
| | PPWD1 | 74 | 31 | 20 | Not modelled | -/SPAC10F6.03c |
| | PPIG | 89 | - | 1 | Not modelled | -/SPAC22H10.11c |
| Substrate | Exon | | | | /E:-12-11 | |
| | Intron | | | | /I:1-23,84-104,128-133 | |

[a] Tandem MS/MS was performed on P complexes stalled with hPrp22 K594A.

[b] Shown in ChimeraX selection notation, where / indicates chain ID and : indicates residue IDs, e.g. /2:4-47 is chain 2 (U2snRNA) residues 4-47.
